# ATPLyzer – An advanced ratiometric multi-colour biosensor for long-term monitoring of ATP dynamics

**DOI:** 10.64898/2026.03.14.711787

**Authors:** Athanasios Papadopoulos, Christian F. Kaiser, Patrick Schlumpberger, Jacqueline Eßer, Jens Reiners, Christoph G.W. Gertzen, Guido Grossmann, Sander H.J. Smits

**Affiliations:** Center for Structural Studies, Faculty of Mathematics and Natural Sciences, Heinrich Heine University Düsseldorf, Düsseldorf, Germany; Cell and Interaction Biology, Faculty of Mathematics and Natural Sciences, Heinrich Heine University Düsseldorf, Düsseldorf, Germany; Institute for Pharmaceutical and Medicinal Chemistry, Faculty of Mathematics and Natural Sciences, Heinrich Heine University, Düsseldorf, Düsseldorf, Germany; Institute for Biochemistry, Faculty of Mathematics and Natural Sciences, Heinrich Heine University Düsseldorf, Düsseldorf, Germany

**Author notes:** **Corresponding Author Sander H.J. Smits** - Center for Structural Studies, Faculty of Mathematics and Natural Sciences, Heinrich Heine University Düsseldorf, Düsseldorf, Germany and Institute for Biochemistry, Faculty of Mathematics and Natural Sciences, Heinrich Heine University Düsseldorf,40225 Düsseldorf, Germany.

## Abstract

Adenosine triphosphate (ATP) is a central molecule in cellular metabolism, serving as the primary energy currency that links catabolic and anabolic pathways. Monitoring intracellular ATP *in vivo* is essential for understanding the dynamics of metabolic states, as well as intracellular functions and intercellular interactions in health and disease. We report the design and application of ATPLyzer, a series of genetically encoded, ratiometric biosensors for the monitoring of ATP levels in living cells. The matryoshka design consists of an ATP-binding cassette linked to a circularly permutated GFP coupled with an internal large stokes shift reference fluorophore, allowing for single-wavelength excitation and ratiometric output. This design overcomes limitations of conventional biosensors, reliance on dual excitation wavelengths, and susceptibility to photobleaching. Multi-colour ATPLyzer variants with different dissociation constants were characterized *in vitro*, exhibiting high specificity for ATP over ADP. Monitoring ATP in *Escherichia coli* confirmed *in vivo* utility and revealed growth-phase and carbon-supply-dependent ATP dynamics. The ATPLyzer biosensor offers a robust and tuneable tool for minimally invasive, time-resolved monitoring of intracellular ATP dynamics.

## Introduction

Adenosine triphosphate (ATP) is fundamental to cellular life, acting as the primary energy currency and driving critical biological processes such as active transport, protein synthesis, intracellular signalling, and metabolic regulation. It also serves as a signalling molecule, mediating intercellular communication, modulating ion channel activity, and regulating key metabolic pathways^1^. Monitoring the tight spatiotemporal regulation of ATP levels is crucial for our understanding of cellular homeostasis, tissue function, and organismal interactions such as symbiosis and infection. Furthermore, imbalances in ATP concentrations have been linked to pathological conditions such as cancer, neurodegeneration, and immune dysfunction^2, 3^. These multifaceted roles of ATP underscore the need for advanced tools to monitor its concentration, distribution, and dynamics in living systems.^4^

Genetically encoded biosensors have emerged as powerfull tools for real-time visualization of ATP dynamics with high spatial and temporal resolution^5–8^. Among the vast types of ATP biosensors, circularly permuted fluorescent protein (cpFP)-based biosensors and Förster resonance energy transfer (FRET)-based biosensors represent the most widely used and impactful classes^6, 8–10^. The cpFP-based biosensors integrate a circularly permuted fluorescent protein into a ligand-binding protein or domain, enabling direct modulation of fluorescence upon ligand binding^6, 9, 10^. Thus, cpFP-based biosensors are proving to be superior due to their unique design and functionality, showing high sensitivity and high signal-to-noise ratio^5, 9, 10^. Numerous cpFP-based biosensors have been developed for different ligands, including the most prominent groupof G-CaMP biosensors for visualization of intracellular Calcium dynamics^11–15^. Notable examples for ATP-specific biosensors include Perceval, which monitors the ATP-to-ADP ratio by exploiting nucleotide-induced conformational changes in the bacterial regulatory protein GlnK1^16^ and ATPQueen, a highly specific cpFP-based biosensor for ATP with increased sensitivity and minimized interference from ADP^9^. Perceval and ATPQueen have been instrumental in elucidating rapid ATP turnover during oxidative stress and metabolic dysfunction^9, 16^.

In comparison, FRET-based ATP biosensors, such as the widely used ATeam, couple the ATP-binding domains to two fluorescent proteins that generate a FRET signal in response to ATP binding^8^. Multiple ATeam variants have been optimized for various affinity ranges and applied across diverse cell types and subcellular compartments, such as mitochondria, the cytoplasm, and the nucleus^8, 17, 18^. Although these biosensors provide ratiometric readouts, which are well-suited for quantitative measurements of ATP gradients in tissues and provide critical insights into ATP gradients in neuronal activity and synaptic function, the rely on FRET between a donor and acceptor and do not support a direct fluorescent correlation to ATP binding^8, 17^.

Despite their utility, cpFP-based biosensors offer significant advantages over FRET-based biosensor systmes. By coupling fluorescence signals to direct metabolite binding events due to conformational modulation of the reporter FP by the ligand-binding protein, they enable more direct detection of ATP levels^5, 9, 10, 19–21^. In fact, cpFP-based biosensors rely on a single fluorophore, making them less prone to photobleaching, simplyfing imaging, and often result in improved signal-to-noise ratios as well as faster kinetics^10, 20, 21^. In contrast, while FRET-based biosensors like ATeam excel in qualitative precision and offer broader dynamic ranges, their dual-fluorophore design introduces complexity and higher susceptibility to experimental artifacts, such as spectral overlap and photobleaching over extended imaging periods^21^. Consequently, the slower response kinetics could limit their application in scenarios requiring rapid detection of ATP flux. However, these limitations highlight the growing preference for cpFP-based biosensors in studies requiring high sensitivity, fast kinetics, and reduced experimental complexity^5, 21^. In turn, the lack of a second FP in the single cpFP-based biosensors is not benefitial for referencing and does not completely support qualitative analysis. Therefore, to overcome the limitations given by the FRET-based and single-FP biosensors, the Matryoshka design has been introduced to create robust genetically encoded biosensors that function ratiometrically^22^. The distinctive molecular architecture of this biosensor design is also reflected in its name: Matryoshka, just like the traditional nesting dolls, the biosensor cassette here contains a reference FP that is nested within the synthetic linker region of the circular-permutated reporter FP constituting the biosensor cassette, which is also nested in the ligand-binding protein^22^. Here, the reference FP is a large stokes shift FP, whereby its excitation wavelength is spectrally far separated (~100 nm) from its emission wavelength^22^. The non-responsive reference FP is capable of being excited by the same wavelength as the responsive FP, allowing for simultaneous imaging, thus reducing the energy intake into the specimen, and essentially doubling the time resolution as compared to FRET-based ratiometric biosensors^22, 23^.

In this study, we present ATPLyzer an advanced ratiometric Matryoshka biosensor that monitors intracellular ATP levels. We were able to re-design a pre-existing intensiometric biosensing module into a ratiometric fashion that tracks ATP levels in the same range as previously reported for other biosensors^9^. Furthermore, our biosensor allowed long-term monitoring of ATP *in vivo*. Taken together, the re-design and creation of ATPLyzer simplified the applicability for ATP biosensors and enables the robust detection of dynamic ATP levels in living systems.

## Results and Discussion

### Molecular design of a Matryoshka-like biosensor for ATP

The bacterial FoF1-ATP synthase represents an ATP:ADP regulated nanomachine with multiple subunits. The ε-subunit of FoF1-ATP synthase has been used priorly for the creation of genetically encoded fluorescence-based biosensors for ATP, representing a suitable recognition element for ATP^8–10^. In aim to design a ratiometric biosensor for ATP, that is not based on FRET, the Matryoshka design provided an opportunity for elevated applicability and simplicity^22, 23^.

To this end, we created the ATPLyzer biosensor family, linking the FoF1-ATP synthase ε-subunit to a responsive FP with a nested, large stokes shift (LSS-)FP as intramolecular reference. LSS-FPs were inserted into the synthetic linker region (GGS-GGT) of a circular permuted GFP (cpGFP) (Figure 1). These nested incorporations of the LSS-FPs into the same molecular cassette with the reporter cpFP enabled the creation of more powerful biosensors with single-excitation wavelengths for both FPs (Figure 1). Furthermore, the usage of two different LSS-FPs (LSSmOrange, Figure 1, A and LSSmApple, Figure 1, B) allows for an optimized and tunable signal-to-noise ratio for a broad range of applications. To creat novel ratiometric biosensors for ATP, the ε-subunit of FoF1-ATP synthase of Bacillus subtilis and PS3 were utilized as described previously^9^. This allowed the creation of three biosensor variants, aming milimolar and micromolar affinities for ATP as well as a non-responsive mutant.

**Figure 1:**
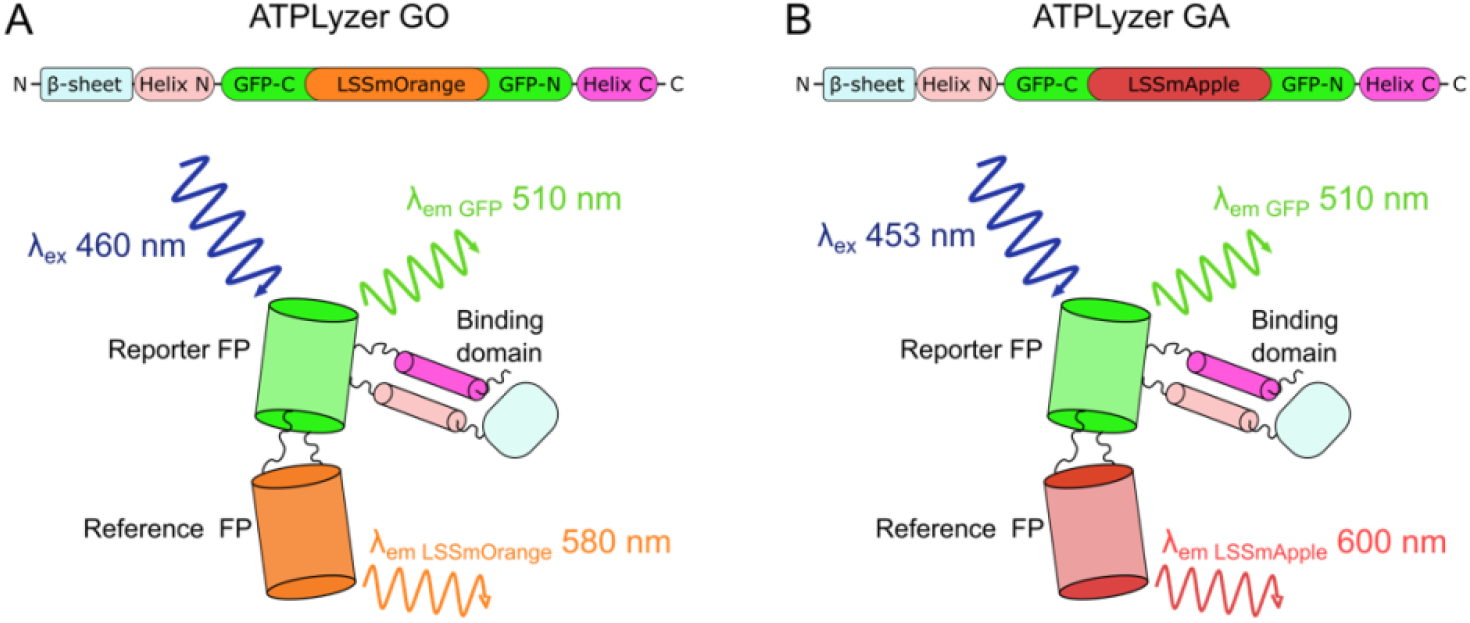
ATPLyzer – a multi-colour biosensor for ATP. The schematic representations of ATPLyzer multi-colour variants are shown for two biosensor cassettes with (**A**) cpGFP-LSSmOrange and ((B)) cpGFP-LSSmApple. The binding domain constitutes of the ε-subunit of the bacterial FoF1-ATPase (β-sheet, Helix N and Helix C). The excitation (λex) and emission wavelengths (λem) are indicated. The colouring corresponds to the parts in the biosensor cassette depiction.

### Characterization of ATPLyzer – a Matryoshka-like biosensor for ATP

Motivated by previous ATP biosensors, three biosensor variants were created aiming for varying affinities in the millimolar and micromolar range as well as a non-responsive mutant^9^. To investigate the binding characteristics of ATPLyzer, the three created biosensor variants were succesfully expressed and purified under identical conditions to ensure comparability of their binding properties (Figure S1). All ATPLyzer variants were generated with a circular permutated GFP as the reporter FP and either LSSmOrange (GO) or LSSmApple (GA) reference FPs. That led to multi-color variants of novel Matryoshka-type biosensors which were investiageted by *in vitro* spectrofluorometry upon titration of ATP and ADP (Figure 2). The fluorimetric analysis of ATPLyzer showed an inverse mode of sensing action, whereby the fluorescence emission intensity of the reporter FP (cpGFP) decreases with increasing ATP concentration, while the emission of reference FPs remains unchanged (Figure 2, A-F). The ATPLyzer GO variants indicated an increased fluorescence intensity for the reference FP with a peak maximum at 575 nm in comparison to the reporter FP (Figure 2, A-C). In turn, the ATPLyzer GA variants indicated moderate levels of fluorescence intensity for the reference FP, indicating a superior signal-to-noise proportion (Figure 2, D-E). While the titration of ATP resulted in a decreased fluorescence intensity of the reporter FP at 510 nm (Figure 2, A,B and D,E), no such effect was observed upon titration of ADP (Figure S2) indicating ATPLyzer’s specificity for ATP. In addition, also no response in terms of fluorescence intensity changes of the reporter FP was observed for the non-responsive (n.r.) variants of ATPLyzer GO and GA, being in line with the expectations (Figure 2, C,F and Figure S2).

**Figure 2:**
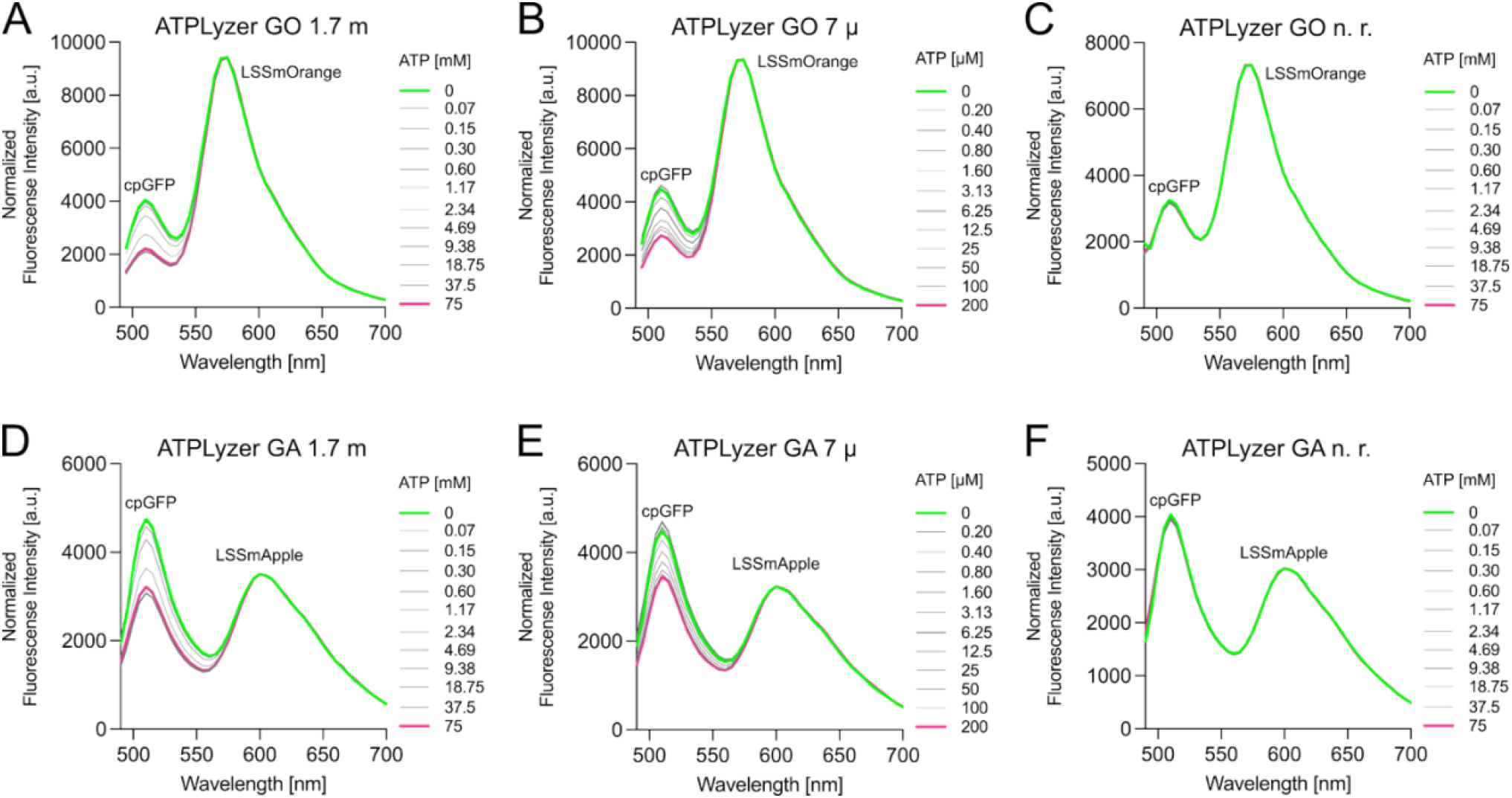
*In vitro* fluorimetric characterization of ATPLyzer. Multi-colour variants of ATPLyzer with varying reference fluorophores (LSSmOrange and LSSmApple) were used for titration of ATP. ATPLyzer variants are shown for two biosensor cassettes with cpsfGFP-LSSmOrange (GO, A, B, C) and cpsfGFP-LSSmApple (GA, D, E, F). For the mM-affinity and non-responding (n.r.) variants of ATPLyzer (A, B 1.7 m and E, F n.r.) 0-75 mM and for ATPLyzer 7 µ (C, D) variant 0-200 µM ATP were used for titration analysis. The experiment conducted with purified ATPLyzer variants and mean values of three biological replicates (n=3) are presented.

Investigations on nucleotide binding revealed that the ATPLyzer variants with lower affinity towards ATP exhibited K_*D*_’s of 1.73 ± 0.04 mM for GO and 1.76 ± 0.06 mM for GA (Figure 3, A and B) and were named as ATPLyzer 1.7 m. The higher affinity variants of ATPLyzer indicated K_*D*_’s of 6.98 ± 0.05 µM for GO and 11.45 ± 1.09 µM for GA (Figure 2, C and D) and named as ATPLyzer 7µ. Neither variant indicated binding towards ADP (Figure S2 and 3). The ATPLyzer n. r. variants did not indicate binding of ATP or ADP upon titration of these nucleotides (Figure S2 and Figure 3) The results are in line with observations made for pre-existing ATP biosensors^9^. To further characterize and elucidate the putative mode of action of ATPlyzer in solution structural analysis was conducted. The proteins appeared monomeric in-solution as indicated by structural analysis via small-angle X-ray scattering (SAXS) (Figure 4 and Figure S4 and S5). Initial AlphaFold3 models of the ATPLyzer Biosensor showed only a low agreement with the collected experimental scattering data (χ^2^ values >5) (Figure S4 and S5). Therefore, we created a library of different conformations and scored the models against the collected experimental data set to identify the in-solution behavior of ATPLyzer. In the apo state the structure of the biosensor is more relaxed and the helices of the FoF1-ATPase ε-subunit (εFoF1) maintain flexibility and freedom to move apart as indicated in the structure (Figure 4, A). When ATP is bound on side of the FoF1-ATPase ε-subunit the helices are trapped in a more compact and static conformation as indicated by the ATP-bound structure (Figure 4, A). The coordination and/or binding of ATP is due to interactions with particular arginines on facing outwards of the εFoF1 helices (Figure 4, B). Docking ATP and ADP to these structures resulted in no interactions of ADP with the C-terminal helices, yet ATP bound in a way that connected the helices but is different from published binding modes. While this is an indicator for the inability to sense ADP, judging from published binding modes, e.g., from the X-ray crystal structure 2E5Y^24^ suggesting ATP-regulated arm motion of its C-terminal domain in F1, ADP cannot bridge the positively charged arginines R92, R99, and R122 using its negatively charged phosphate groups, compared to ATP (Figure 4, C and D).These results are in line with previous research and provide evident data on structural arrangements that were proposed for the mode of action of the ε-subunit of FoF1-ATP synthase^9, 10, 24^.

**Figure 3:**
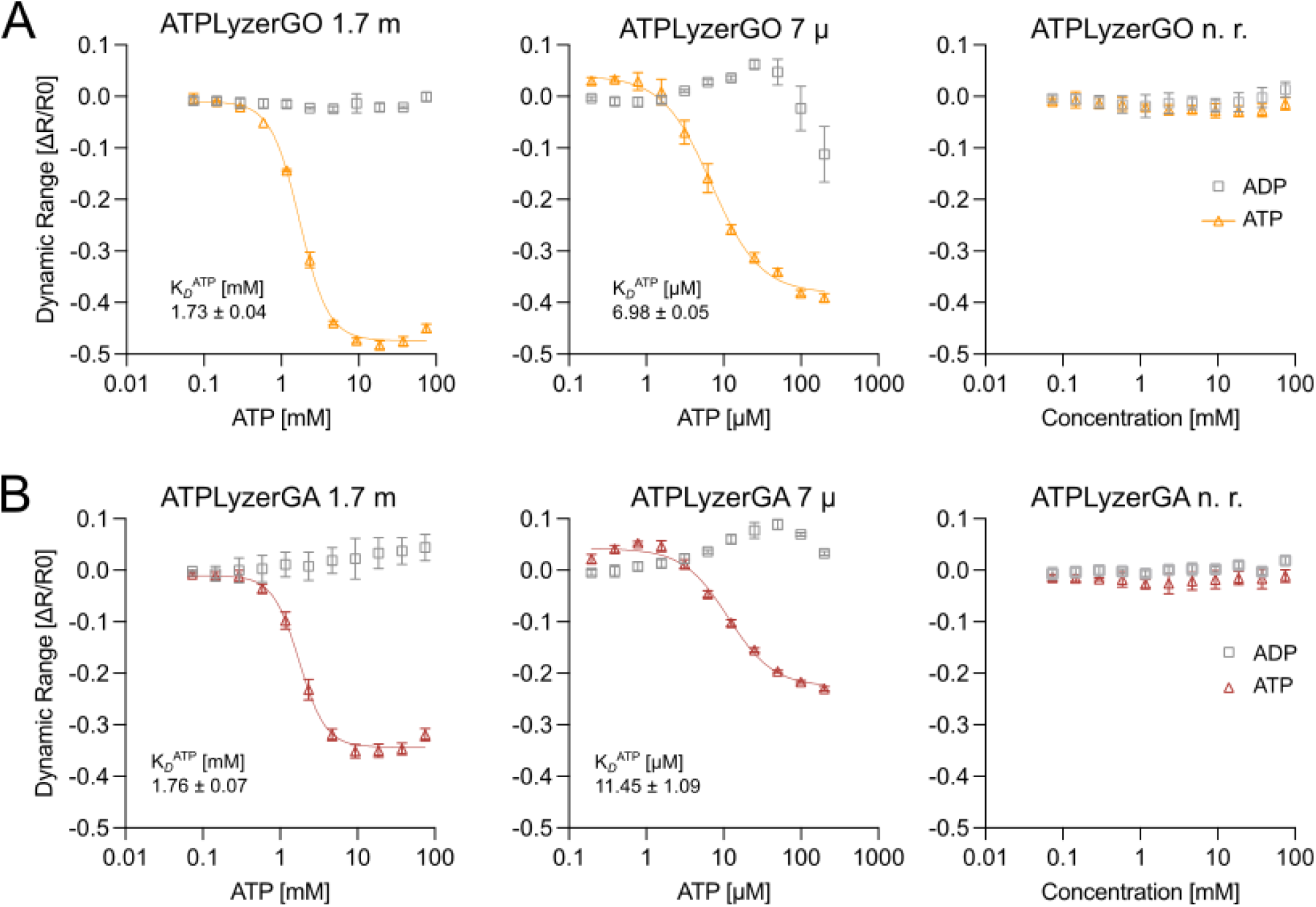
*In vitro* nucleotides binding by ATPLyzer. Multi-colour variants of ATPLyzer with varying reference fluorophores (LSSmOrange and LSSmApple) were used for titration of ATP and ADP. ATPLyzer variants are shown for two biosensor cassettes with (**A**) cpGFP-LSSmOrange (GO) and (**B**) cpGFP-LSSmApple (GA). For the mM-affinity and non-responding (n.r.) variants of ATPLyzer (1.7 m and n.r.) 0-75 mM and for ATPLyzer 7 µ variant 0-200 µM ATP were used for titration analysis. The dissociation constant (Kd’s) The experiment conducted with purified ATPLyzer variants and mean values and the standard deviation (SD) of three biological replicates (n=3) are presented.

**Figure 4:**
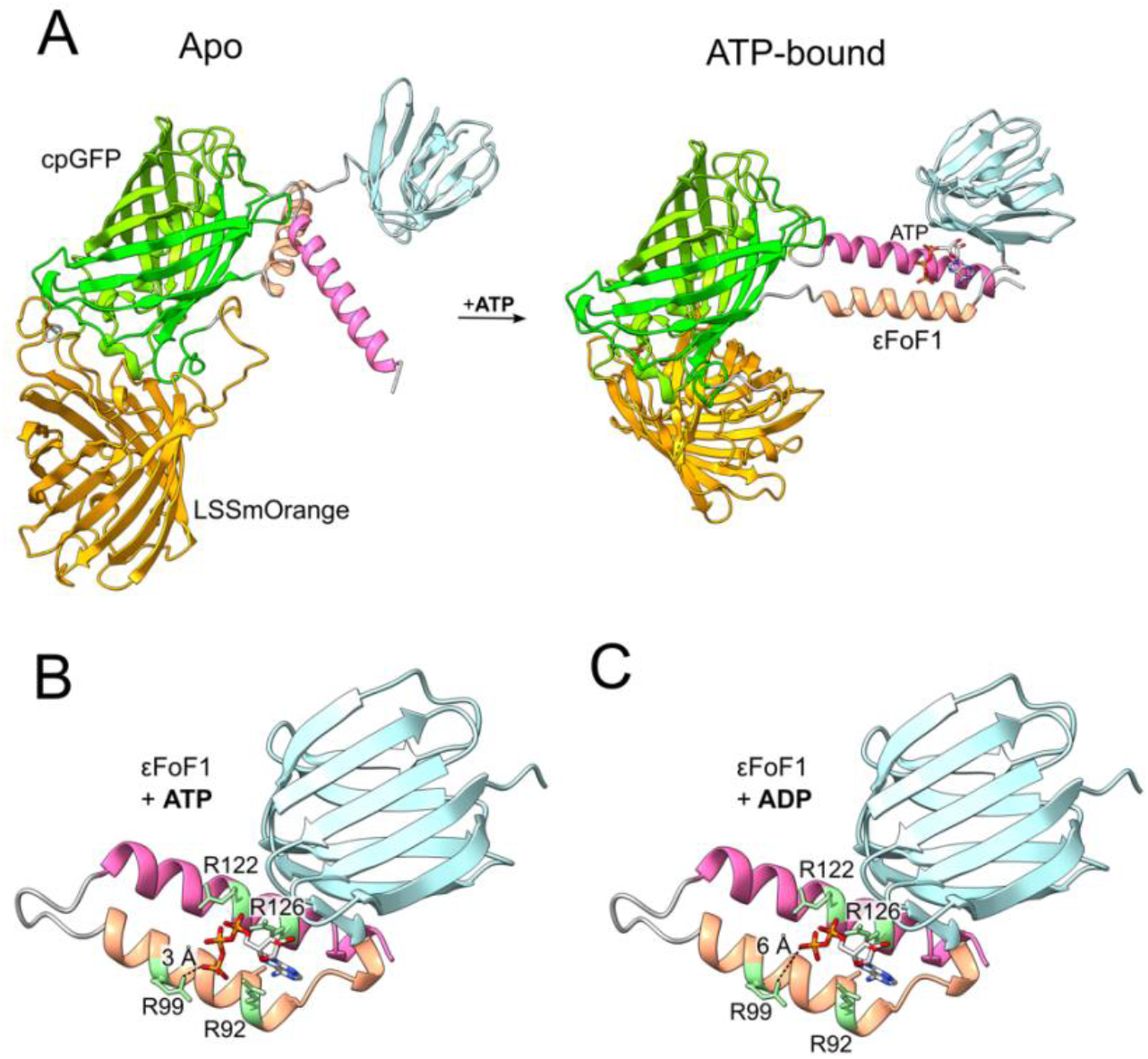
Structural analysis of ATPLyzer. ATPLyzer green-orange (GO) variant 1.7 m in solution structural behavior by SAXS. The reporter FP (cpGFP) in green, the reference FP (LSSmOrange) in orange. The FoF1-ATPase ε-subunit (εFoF1) forms the molecular binding element of the biosensor and compose of the N-terminal β-sheet in turquoise, the N-terminal helix in salmon, the C-terminal helix in pink. (**A**) In the Apo form the structure the Biosensor is more relaxed and the helices of the FoF1-ATPase ε-subunit (εFoF1) maintain freedom to move as indicated by large separation in the structure. When ATP is bound on side of the FoF1-ATPase ε-subunit the helices are trapped in a more rigid conformation as indicated by the „ATP-bound” structure. The coordination and binding of ATP is due to interactions with particular arginines on facing outwards of the εFoF1 helices. Further SAXS data can be found in the supplementary information. Binding mode comparison of ATP (**B**) and ADP (**C**) in the X-ray crystal structure of the thermophilic bacillus PS3 (PDB-ID: 2E5Y) showing that ATP can bridge the arginines (R92 and R99) in the N-terminal helix and R122 and R126 in the C-terminal helix, while ADP cannot as indicated by the distance indicated to R99 (Å). The models are tilt by 30° to the front in comparison to (A) for complete view on nucleotide coordination to εFoF1.

### *In vivo* expression of the ATPLyzer reveals growth-dependent ATP levels in *Escherichia coli* populations

To evaluate whether ATPLyzer 7µ is functional in living cells, we assessed its performance in the gram-negative model *Escherichia coli* XL1-Blue, part of the K-12 lineage, (*E. coli*) in a multi-well reader experiment. The nested reference fluorophore design of the biosensor enabled hereby the averaging of population level ATP-dependent fluorescence shifts. Both color variants of ATPLyzer, GO and GA, were utilized for investigations (Figure 5). Therefore, the active biosensor, the non-responsive (n.r.) variant and wild-type *E. coli* strain were chosen to validate the observed fluorescence dynamics and ATP metabolism. ATP-dependent changes were expected to be solely detectable in the active biosensor and responses to carbon supplementation and perturbations of ATP synthesis. Mid-exponential (OD_600nm_ 0.5-0.8) cultures, grown in rich LB medium, were transferred to minimal carbon-free M9 medium initiating a transient starvation period of 3 h. Glucose was then supplemented to allow growth and followed by a growth period of 23.5 h. In the stationary phase, the protonophore carbonyl cyanide 3-chlorophenylhydrazone (CCCP) (Figure 5) was then supplemented to dissipate the proton motive force (PMF) and inhibit ATP synthesis^25–28^, providing a validation of *in vivo* responsiveness. As indicated by the monitoring of the optical density (OD600) of the cultures, no growth defects were observed due to the biosensor expression in comparison to the control *E. coli* wild-type strain (Figure 5, OD_600_).

**Figure 5:**
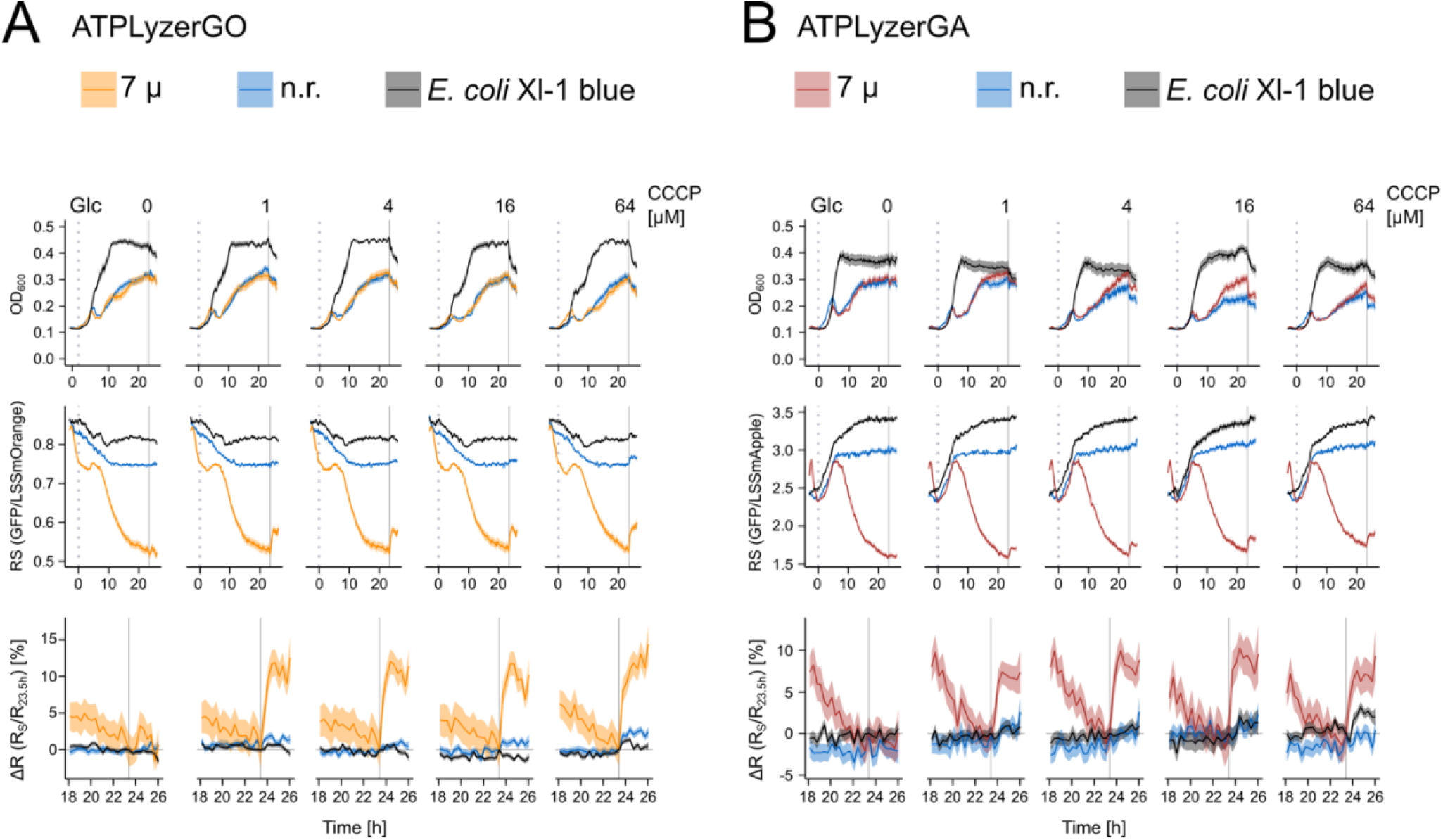
*In vivo* application ATPLyzer in bacterial cells. *E. coli* XL1-blue populations expressing the ATPLyzer with (**A**) LSSmOrange or (**B**) LSSmApple reference fluorophores were grown in M9 medium, starved for 3 h, and supplied with 625 µM glucose (Glc, grey dashed line), followed by a single CCCP (grey solid line, t = 23.5 h) treatment of 0-64 µM in the stationary phase. Cultivation was performed in 96-well plates with automated measurements every 210 s, and values were averaged over 20-min intervals. First plots indicate the optical density of the cell culture measured at 600 nm (OD_600_) over time. The middle panels indicate the course of biosensor response Rs (GFP/reference ratio) upon glucose and CCCP addition. The third panels depict a zoomed-in view of Rs dynamics relative to the last non-CCCP-treated timepoint at 23.5h after CCCP addition at different concentrations. Shaded regions indicate mean ± SEM. Means were calculated from three independent experiments with four wells per condition (N = 12 total measurements per condition).

The initial starving period in the ATPLyzer 7µ display a transient increase of the ratiometric response (Figure 5, RS) indicating a short-timed decrease of ATP-binding, followed by a consistent decrease of the ratio until glucose was supplemented. After the addition of glucose growth was regained within 2 h, while the RS continuously rose until a peak at 6 h post treatment, which was followed by a progressive decline in RS gradually stabilizing to a lower plateau. Addition of CCCP at 23.5 h displayed a rapid increase in ratio declining post-treatment, matching a decline in ATP production associated with expected disruption of the proton motive force. The CCCP-dependent signal change ΔR_s_ was calculated against the last timepoint before treatment (23.5 h) displaying a 10-15 % increase of the signal for the active sensor variant. In contrast, neither the wild-type *E. coli* nor the n.r. variant responded with a similar shift in ratio to CCCP treatment. Further, the autofluorescence and n.r. variant derived ratio patterns deviated between the chosen emission wavelengths with LSSmOrange (Figure 5, A/RS) displaying a negative trend throughout growth, while LSSmApple (Figure 5, B/RS) displayed a positive trend. Both controls displayed a progressive increase or decrease of the ratio values, dependent on the emission wavelength captured, which gradually stabilized in the stationary phase. Notably the n.r. variant remained steadily within the wildtype and active biosensor variant, leading to a clear difference between the active biosensor, n.r. variant and wildtype cultures.

To distinguish metabolic activity from carbon presence, we compared glucose with non-metabolizable sucrose in *E. coli* K-12 lacking csc genes, and restored sucrose utilization by adding invertase (Figure S3). Sucrose alone did not support growth, whereas invertase restored growth similar to glucose. Metabolizable carbon induced a transient signal flattening followed by a peak, delayed at higher concentrations and more pronounced in the LSSmApple variant. Sucrose alone resembled the 0 mM control, showing a continuous decline.

Together, these results highlight the *in vivo* responsiveness of the ATPLyzer 7µ reflecting negative and positive perturbations of ATP production dynamically and across a population of live bacterial cells. Our results reveal transient shifts in ATP availability by the biosensor following disruption of the proton motive force, starvation and growth-dependent shifts.

## Conclusion

This study presents the development and comparative biochemical analysis of a novel genetically encoded ratiometric biosensor for adenosine triphosphate (ATP), designed to overcome limitations of previously existing fluorescent biosensors for ATP^9^. The biosensor, termed ATPLyzer, builds upon the architecture of the pre-existing single-FP ATP biosensors and incorporates a nested (Matryoshka-like) dual-fluorophore design, enabling full ratiometric measurements with enhanced sensitivity and stability (Figure 1)^22^. These established benchmarks served as the foundation for engineering the ATPLyzer biosensor. The novel and refined design of ATPLyzer demonstrated clear advantages in signal consistency, operational robustness, and fluorescence intensity, even under variable intracellular and extracellular conditions (Figure 2). A critical advancement of ATPLyzer lies in its ability to perform consistent ATP detection irrespective of fluctuating expression levels, due to its ratiometric output, which normalizes signal to a stable reference fluorophore. This represents a significant improvement over intensity-based biosensors like ATPQueen, which can be confounded by differences in protein expression or photobleaching ^9^. Additionally, the capability of multi-colour design of ATPLyzers biosensor cassettes, either with green-orange (GO) or green-aoole (GA) fluorescence emission allows for broad range of utilizations and adaptability to the requirements of the analysed living systems. Biochemical analysis revealed distinct performance characteristics of the ATPLyzer biosensor variants, which stem from their divergent protein domains derived from different *Bacillus* species (Figure 3 and 4). Unlike prior biosensors, ATPLyzer variants maintain functionality across a broad range of ATP concentrations and buffer conditions. Furthermore, the biosensor function is not limited to ambient temperature as indicated best for previous biosensors^9^ but also works at common bacterial cultivation conditions, including 37°C. The present results show the capability of ATPLyzer 7µ to report on *in vivo* ATP dynamics in *E. coli* populations. Both ATPLyzer GO and GA biosensor variants tracked similarly CCCP-mediated inhibition of ATP synthesis, glucose-driven growth recovery and starvation periods (Figure 5). Comparisons of glucose with non-metabolizable sucrose, together with invertase rescue, confirmed that ATPLyzer output depends on carbon catabolism rather than carbon presence alone (Figure S3). The ATPLyzer 7 µ variants exhibited reliable ATP-induced fluorescence modulation *in vitro* and *in vivo*, while the 1.7 m variants displayed brought capability *in vitro*. These findings underscore the tunability of the ATPLyzer biosensors based on the ε-subunit employed, enabling tailored application across diverse cellular environments with varying ATP levels and ionic compositions. Furthermore, the ratiometric nature of ATPLyzer eliminates the need for extensive normalization procedures and facilitates more accurate quantification of ATP dynamics in living cells. This advantage becomes particularly relevant in high-throughput or longitudinal imaging applications where photostability and expression variability can obscure subtle changes in analyte concentration.

Taken together, the ATPLyzer biosensor represents a significant improvement over previous ATP biosensors, offering specificity for ATP with tuneable affinities and enabling true ratiometric output, thereby providing a powerful tool for probing ATP-mediated cellular processes with greater precision and reliability.

## Material and Methods

### Construction of matryoshka biosensor for ATP

To construct ATPLyzer, the gene of the large-stokes-shift fluorophore LSSmOrange was integrated into pre-existing ATPQueen^9^ gene cassette decoded in pRSET_B_-based plasmid^1^. The molecular cloning was conducted by Gibson assembly (NEB) based on the manufacturer’s guidelines.

### Expression and purification of the biosensor

For the expression of biosensors, the corresponding plasmids were used for transformation of chemically competent *E. coli* BL21 (DE3) by heat-shock method^6^. In brief, 50 ng/µl of the plasmids were added to 100 µl cell solution and incubated on ice for 15 min, following a 45 second heat-shock at 42°C with subsequent incubation on ice for 3 min. Afterwards, 500 µl lysogeny broth (LB, Luria/Miler, Roth) were added to the bacterial solution and mixed by pipetting. The solution was transferred to LB agar plates containing 100 µg/ml ampicillin. The plates were incubated at 37°C overnight and stored at 6°C until further usage. Positive transformants were identified by their fluorescence properties using a commercial fluorescence imaging (Amersham IQ800 Imager, GE). To initiate the cultivation of positive transformants and subsequent heterologous protein expression, 10 ml of LB medium supplemented with ampicillin (100 µg/ml) was dispensed into a sterile 20 ml reagent tube under aseptic conditions. Fluorescent bacterial colonies indicative of successful transformation was identified using a fluorescence imaging system (Amersham IQ800, GE Healthcare) and were selected for inoculation of pre-expression cultures.

The inoculated cultures were incubated at 37°C with agitation at 220 rpm in darkened conditions to minimize photobleaching of fluorophores. Bacterial growth was monitored by measuring the optical density at 600 nm (OD_600_) at regular intervals. Upon reaching mid-logarithmic phase (OD_600_ ≈ 0.6), 1.5 ml of the pre-culture was used to inoculate 100 ml of auto-inductive (AIM) expression medium^7^. The AI medium, consisting of LB supplemented with 0.2% (w/v) lactose and 0.05% (w/v) glucose, was pre-prepared and dispensed into a 500 ml baffled Erlenmeyer flask containing ampicillin (100 µg/ml) to maintain selective pressure.

The auto-inductive system leverages differential metabolism of carbon sources for auto-induction enabling a metabolic shift that allows for auto-regulated induction of recombinant protein expression without the need for manual addition of IPTG or OD-dependent timing. Protein expression was carried out by incubating cultures at 20°C with shaking at 220 rpm for 48 hours in darkened conditions to prevent fluorophore bleaching. Expressed cells were harvested at 4°C with 4350 g for 20 min. Cell pellets were suspended in 50 mM MOPS pH 7.0, 50 mM KCl, 0.5 mM MgCl_2,_ and optionally supplemented with 0.05 % (w/v) Triton X-100, as described in previous protocols^1^.

The cells were lysed by sonification (Process time 45 seconds, Pulse time ON: 3 seconds, OFF: 8 seconds, with total of utilized energy of 633 J, QSonica Q700). Subsequently the solution was applied for centrifugation at 20000 g, 4°C for 10 min to pellet cell debris. The purification of the respective biosensors conducted via immobilized metal affinity chromatography (IMAC) using NiNTA agarose (Protino, Macherey-Nagel) in chromatography columns (Bio-Rad) that were equilibrated with 50 mM MOPS pH 7.0, 250 mM KCl, 0.5 mM MgCl_2_. The histidine-tagged biosensors were immobilized on the NiNTA beads, washed with 50 mM MOPS pH 7.0, 250 mM KCl, 0.5 mM MgCl_2_, 10 mM imidazole to remove weakly bound contaminants. The biosensors were eluted from the beads with 50 mM MOPS pH 7.0, 50 mM KCl, 0.5 mM MgCl_2_, 300 mM imidazole, 5% glycerol, and stored on ice or in the fridge overnight to allow fluorophore maturation in darkened conditions. The sample purity was determined by SDS-PAGE and in-gel fluorescence (Amersham IQ800, GE) as well as conventional colloidal Coomassie staining (Serva). The protein concentration was determined by UV/Vis spectrometry using a NanoDrop (Thermo Scientific).

### Characterization of the biosensor

To evaluate the biosensor’s sensing mode and metabolite-binding capability, fluorimetric titrations were performed using a plate-reader assay (Infinite M Plex, TECAN). Each titration series comprised 12 concentrations generated by serial dilution from row 12 to row 2, with a zero-concentration control in row 1. First, 100 µL of assay buffer (50 mM MOPS pH 7.0, 200 mM KCl, and 1 mM MgCl_2_) were added to wells in rows 1–11, while 200 µL of the metabolite stock solution were added to row 12. For the ATPLyzer 1.7 m and NA variants, ADP or ATP serial dilutions were prepared in the range of 75 mM to 0 mM; for the ATPLyzer 7µM variants, metabolite concentrations ranged from 200 µM to 0 µM. The biosensor solution was diluted to a final protein concentration of 0.1 mg/mL in assay buffer. Subsequently, 100 µL of the biosensor solution (0.1 mg/mL) were added to each well. For buffer controls, 100 µL of the metabolite titration series were mixed with 100 µL of assay buffer lacking the biosensor. The final volume in each well was 200 µL. Plates were incubated for 15 minutes at room temperature in the dark before measurement. For the preparation of ATP and ADP stocks 50 mM MOPS pH 7.0, 200 mM KCl were used and pH adjusted to meet a pH of 7.0, while 1 mM MgCl_2_ was added to the final stocks of 150 mM and 400 µM before each measurement. Immediately prior the measurement, plates were shaken for 30 seconds to ensure homogeneous mixing. Fluorescence emission spectra were recorded from 490 to 700 nm in 5 nm increments following excitation at 460 nm for cpsfGFP-LSSmOrange and 453 nm for cpsfGFP-LSSmApple biosensors. Buffer autofluorescence was negligible. The reporter fluorescent protein (cpsfGFP) displayed an emission maximum at 505–510 nm, while the reference fluorescent protein emitted at 575 nm for LSSmOrange and 600 nm for LSSmApple. For data analysis, the reference FP intensity was used to normalize each data point within the recorded spectra. The values of emission maxima were used to calculate the dynamic range changes [ΔR/R_0_]^8^ in response to analyte binding by the respective biosensors and to assess the sensing mode utilizing the equitation below as described previously^5, 7, 8^.

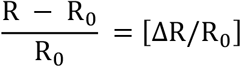

R _0_: Fluorescence emission Intensity ratios (cpsfGFP/REF) prior addition of ligand

R : Fluorescence emission Intensity ratios (cpsfGFP/REF) upon increasing ligand concentrations

The binding affinity was determined by fitting the obtained titration data. The reported Kd values are calculated by the concentration-dependent X values whereby four-parameter logistic (4PL, X is concentration and not log concentration X) was used in a dose-response model to determine the IC50 of the agonist (ferrous iron), which is the concentration that causes a response half-way between the of the “bottom” (basal) and “top” (maximal) response.

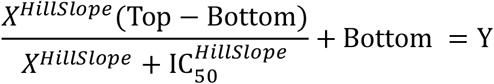

Equitation and supporting information, including the interpret parameters (IC50, Hill Slope, Top and Bottom plateaus of Y axis) given by GraphPad Prism 10.4.0 (527) (GraphPad Software, LLC). The representative IC50 is equal to the Kd values. The error is indicated as standard error of the mean (SEM) obtained by at least three biological replicates (n=3).

### Structural analysis by size-exclusion chromatography coupled small-angle X-ray scattering (SEC-SAXS)

The SEC-SAXS data were collected on beamline BM29 at the ESRF Grenoble ^29, 30^. The BM29 beamline was equipped with a PILATUS 2M detector (Dectris) at a fixed distance of 2.812 m. The measurements were performed at 10°C with a protein concentration of 4.90 mg/ml for the apo protein and for the ATP measurement. The SEC-SAXS runs were performed on a Superdex 200 increase 10/300 column (100 µl inject, 50 mM MOPS pH7.0, 100 mM KCl, 1 mM MgCl_2_ for apo and with additional 5 mM ATP for the ATP measurement) with a flowrate of 0.6 ml/min. A total of 800 frames with an exposer time of 3 sec/frame were collected and scaled the data to absolute intensity against water. All used programs for data processing were part of the ATSAS Software package (Version 3.0.5) ^31^. Primary data reduction were performed with the programs CHROMIXS ^32^ and PRIMUS ^33^. Using the Guinier approximation ^33^, the forward scattering *I(0)* and the radius of gyration (*R* _*g*_) were determined. The program GNOM ^34^ was used to estimate the maximum particle dimension (*D*_*max*_) with the pair-distribution function *p(r)*. The initial models of the ATPLyzer Biosensor were created with AlphaFold3 ^35^. For a better agreement with the experimental data the program RANCH (part of EOM) ^36, 37^ was utilized to create a library of 10000 models, where the linker regions of the ATPLyzer domains as well as the N-terminal tail was set as flexible, while keeping the fluorophores as rigid bodies. The created model library was scored with CRYSOL ^37^ to find a model which is in-line with the experimental data.

### X-ray structure analysis and molecular docking

ADP and ATP were drawn and converted into 3D using the ChemDraw19 suite were docked to the effector-bound structure as modelled via SAXS measurements (*vide supra*) utilizing a combination of AutoDock as a docking engine and the DrugScore2018 distance-dependent pair-potentials as an objective function^38–40^. The ligands were allowed to explore the entirety of their target structures and not confined to the binding pockets. A clustering RMSD cut-off of 3 Å was used and the cluster with the highest docking energy comprising at least 20% of all docking poses was considered valid.

### *In vivo* characterization of the ATPLyzer Biosensor Using Plate Reader Assays

The *in vivo* performance of the ATPLyzer biosensor was tested by fluorescence and absorbance measurements in *E. coli* cultures using the ClarioStar Plus plate reader (BMG Labtech GmbH) operated using the Smart Control software (BMG Labtech GmbH, V.6.3.0) or Voyager (BMG Labtech GmbH, V.2510.1) for CCCP experiments.

All experiments were performed using the *E. coli* strain XL1-Blue (Agilent Technologies, Genotype: *recA1 endA1 gyrA96 thi-1 hsdR17 supE44 relA1 lac [F́ proAB lacIqZΔM15 Tn10* (Tetr)]), a K12 derivate for highly efficient transformation. Cultivation was performed in in lysogeny broth (LB) medium at 37°C and 200 rpm shaking. If applicable ampicillin was supplemented at 100 µg/mL. The non-expressing wildtype of the strain was used as a non-fluorescent control, while genetically modified variants were introduced using heat-shock transformation with the respective plasmid.

Precultures were directly inoculated from the cryo-culture (25 % glycerol, −70 °C) and grown overnight. Main cultures were prepared by inoculating 10 mL of fresh LB medium with 100 µL (OD 4.0) preculture and shaken at 37°C until mid-exponential (OD600 nm 0.5 - 0.8).

Cells were harvested and washed twice with the minimal M9 medium (supplemented with 0.1mM CaCl_2_, 1mM MgSO_4_, 0.05mM FeCl_3_, and 2.52µM trace elements). A final dilution of the washed inoculum in supplemented M9 at an OD600nm of 0.1 was then transferred to the 96-well plate (Greiner Bio One). Peripheral wells were filled with sterile water to reduce edge effects during measurement.

Plates were then measured at 210 s intervals at 37 °C with intermittent 30s 300 rpm double-orbital shaking intervals. Every interval included a measurement of optical density at 600 nm, (470 ± 15 nm / Emission 515 ± 20 nm), and either LSSmApple (490 ± 15 nm / Emission 600 ± 20 nm), or LSSmOrange (437± 15 nm / Emission 572 ± 20 nm). Fluorescence measurement were performed with an automatic gain window adjustment to ensure stable signal output using the “Enhance Dynamic Range” option of the ClarioStar Plus (BMG Labtech GmbH). Carbon sources (20 µL) were added after 3 h of measurement, while CCCP solutions were added after 26 h. In sucrose and invertase experiments, sucrose was added immediately at the start of the experiment and 50 U/mL invertase was added after a 2 h incubation period.

Data from three independent biological replicates were combined and visualized utilizing R in the R-Studio environment^41^. Fully annotated analysis scripts are available in the supplementary materials. The ratiometric response of the sensor in *in vitro* experiment was calculated by averaging all measurements contained within a 20 min interval:

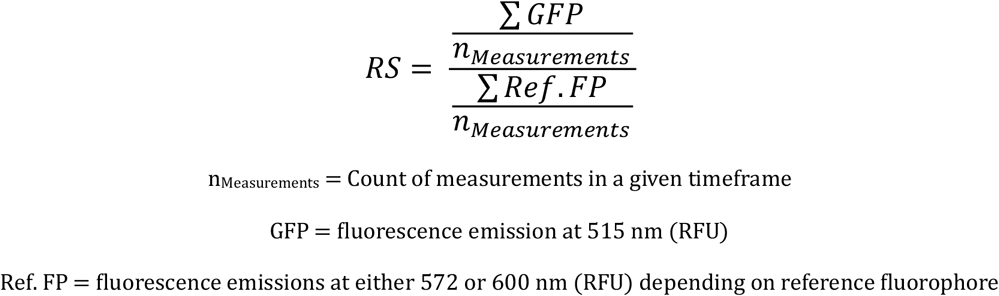

The visualization of CCCP induced effects was performed by calculating a signal change termed ΔR_s_ normalized to the time of treatment in this case t = 23.5 h, resulting in a % change from the before treatment signal:

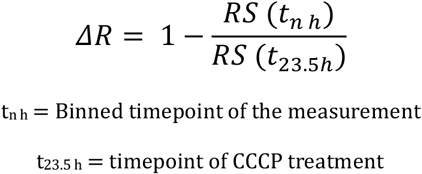

## Supporting information

Supplementary infromation

## Abbreviations

FP: fluorescent protein
FRET: Förster resonance energy transfer
cpFP: circular-permutated fluorophore
sfGFP: super-folder green fluorescent protein
LSS: large stokes shift
Single-FP: single-fluorescent protein or fluorophore
PMF: proton motive force
SEC-SAXS: size-exclusion chromatography coupled small-angle X-ray scattering

## Associated Content

### Data Availability

We uploaded the SAXS data to the Small Angle Scattering Biological Data Bank (SASBDB)^42^, with the accession codes SASDYQ9 (ATPLyzer GO 1.7 m apo) and SASDYR9 (ATPLyzer GO 1.7 m ATP)

## Notes

The authors declare no competing financial interest.

## Author contributions

AP, SS and GG initiated this study, AP conducted the biosensor design, molecular bioengineering, protein characterization. AP, JR and CG conducted the structural analysis. CK, PS and JE conducted *in vivo* experiments. SS and GG coordinated funding project administration and supervised. AP and SS wrote the initial manuscript. All authors contributed to fruitful discussions and lively development of the manuscript. Correspondence and requests for materials should be addressed to Athanasios Papadopoulos and Sander Smits.

## Acknowledgements

Our special thanks to the Institute for Biochemistry for their friendly support in laboratory equipment. We acknowledge the European Synchrotron Radiation Facility (ESRF) for the provision of synchrotron radiation facilities, and we would like to thank Mark Tully for assistance in using beamline BM29. We also acknowledge DESY (Hamburg, Germany), a member of the Helmholtz Association HGF, for the provision of experimental facilities. Parts of this research were carried out at PETRA III, and we would like to thank Cy M. Jeffries and Dmytro Soloviov (EMBL Hamburg) for assistance in using beamline P12. Computational support and infrastructure were provided by the “Zentrum für Informations-und Medientechnologie” (ZIM) at Heinrich Heine University Düsseldorf.

