## Supplementary infromation for "ATPLyzer – An advanced ratiometric multi-colour biosensor for long-term monitoring of ATP dynamics"

\# corresponding author

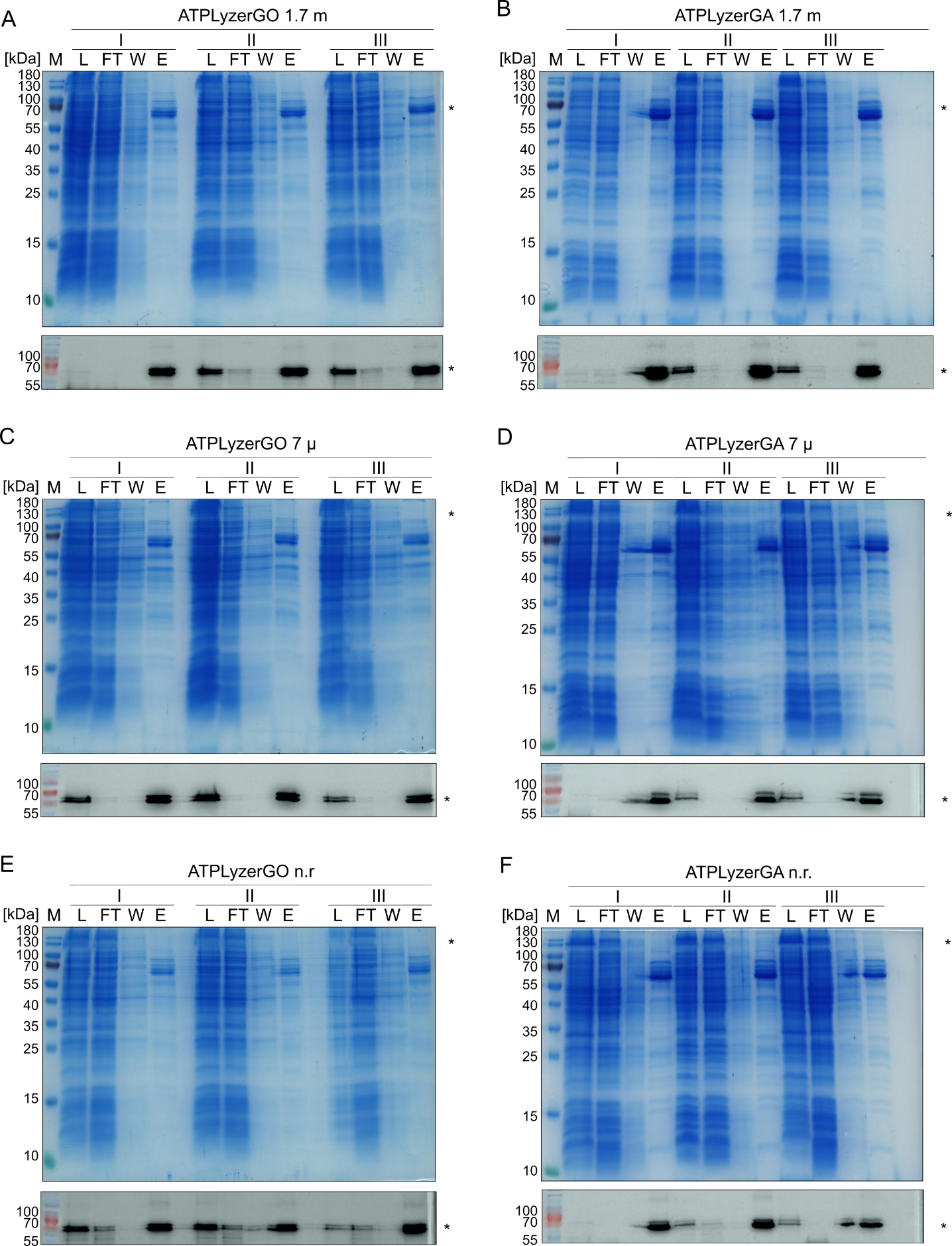

**Figure S1: Purification of ATPLyzer variants.** Multi-colour variants of ATPLyzer with varying reference fluorophores (LSSmOrange ‘GO’ and LSSmApple ’GA’) were heterologously expressed in bacteria and purified by immobilized metal ion chromatography. The SDS-PAGE is shown by Coomassie colloidal stain (above) and *in gel* fluorescence (below). The samples are labelled (A – F) and the analysed corresponding fraction with, L: cell lysate, FT: Flow-through, W: wash fraction, E: elution fraction. The asterisk marks the prominent band of the biosensor. The molecular weight is indicated in kDa (Page Ruler prestained, Thermo Fisher Scientific).

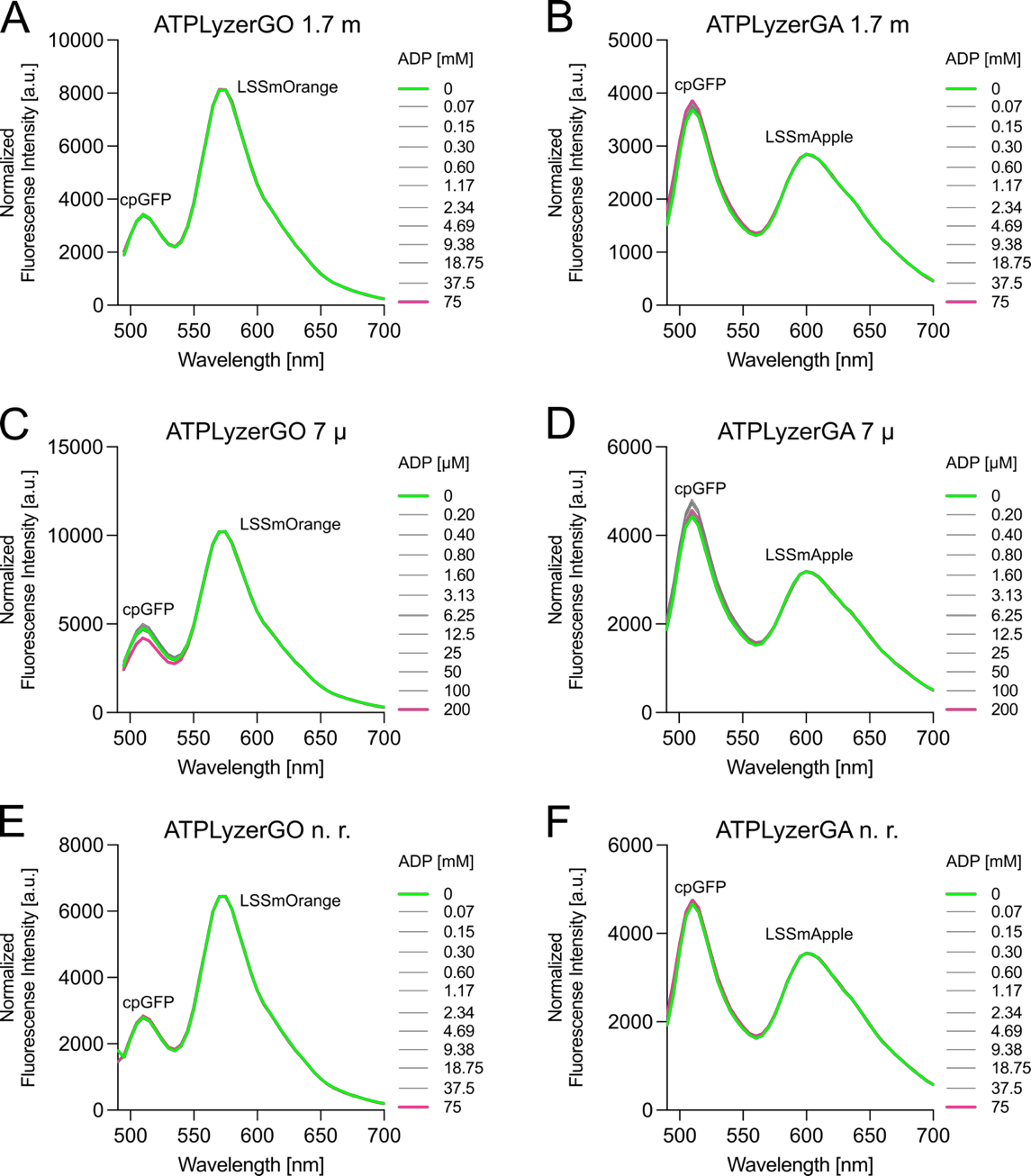

**Figure S2: *In vitro* fluorimetric characterization of ATPLyzer with ADP.** Multi-colour variants of ATPLyzer with varying reference fluorophores (LSSmOrange and LSSmApple) were used for titration of ATP. ATPLyzer variants are shown for two biosensor cassettes with cpsfGFP-LSSmOrange (GO, **A**, **C**, **E**) and cpsfGFP-LSSmApple (GA, **B**, **D**, **F**). For the mM-affinity and non-responding (n.r.) variants of ATPLyzer (**A**, **B** 1.7 m and **E**, **F** n.r.) 0-75 mM and for ATPLyzer 7 µ (**C**, **D**) variant 0-200 µM ADP were used for titration analysis. The experiment conducted with purified ATPLyzer variants and mean values of three biological replicates (n=3) are presented.

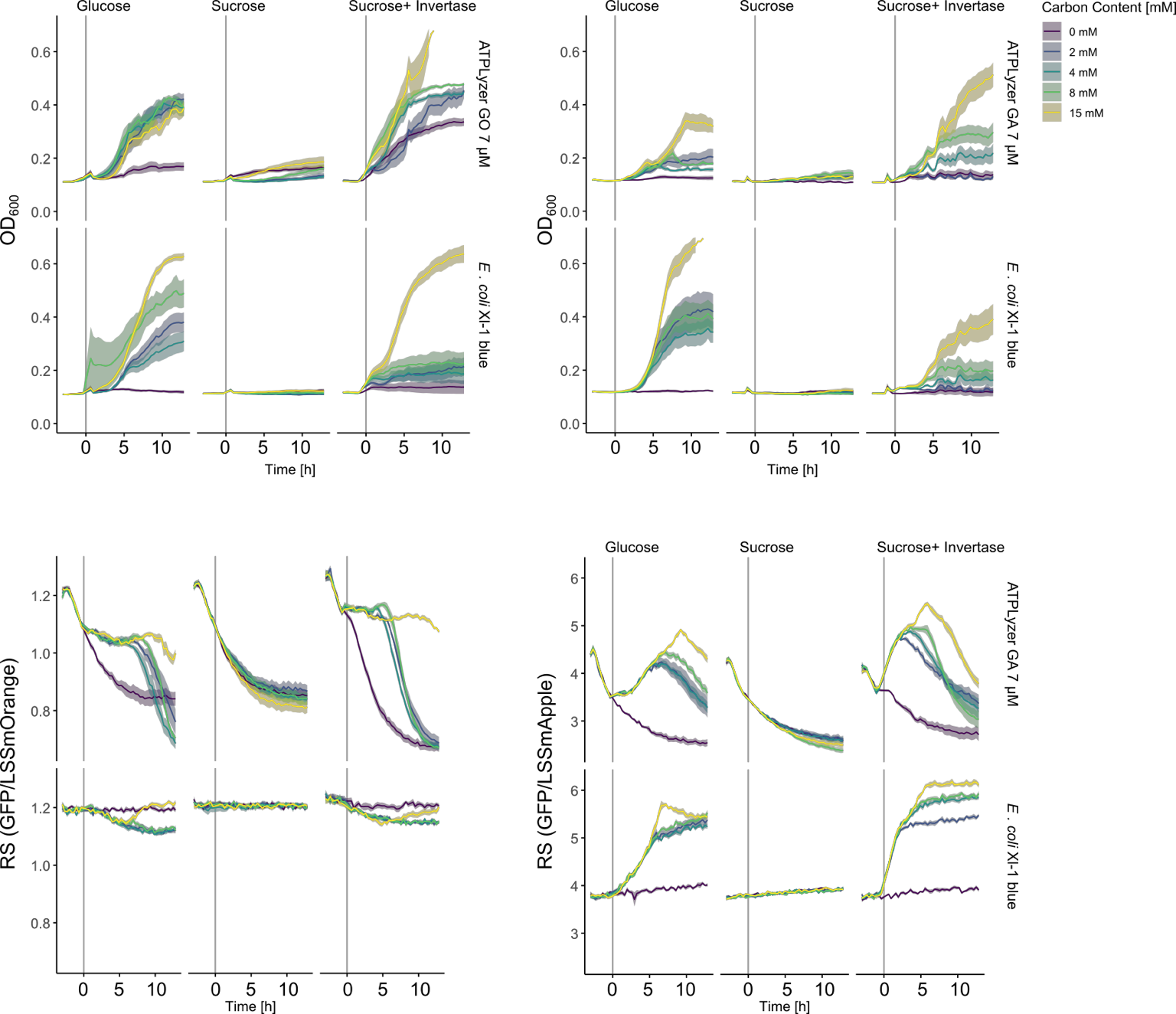

**Figure S3: Exogenous invertase restores ATP-response and growth in sucrose-negative** ***E. coli* XL1Blue.** *csc/scr*-negative *E. coli* XL1-Blue populations expressing the ATPLyzer 7µ with either LSSmOrange (left) or mApple (right). Bacterial cultures were incubated in minimal M9 medium lacking a carbon source, which was supplemented (grey solid line) after 3 hours of starvation (t = 0 h). Carbon was supplemented at various concentrations, as shown in the top right, either in the form of glucose or sucrose, or sucrose supplemented with invertase (10 U/mL). The upper panel displays the cell density as OD_600nm_ and the lower panel the ratiometric sensor signal Rs (FI_GFP_/FI_Ref-FP_). Line plots display the mean signal averaged across x wells captured in x different plates, with the shaded area showing the SEM (N = 18-36).

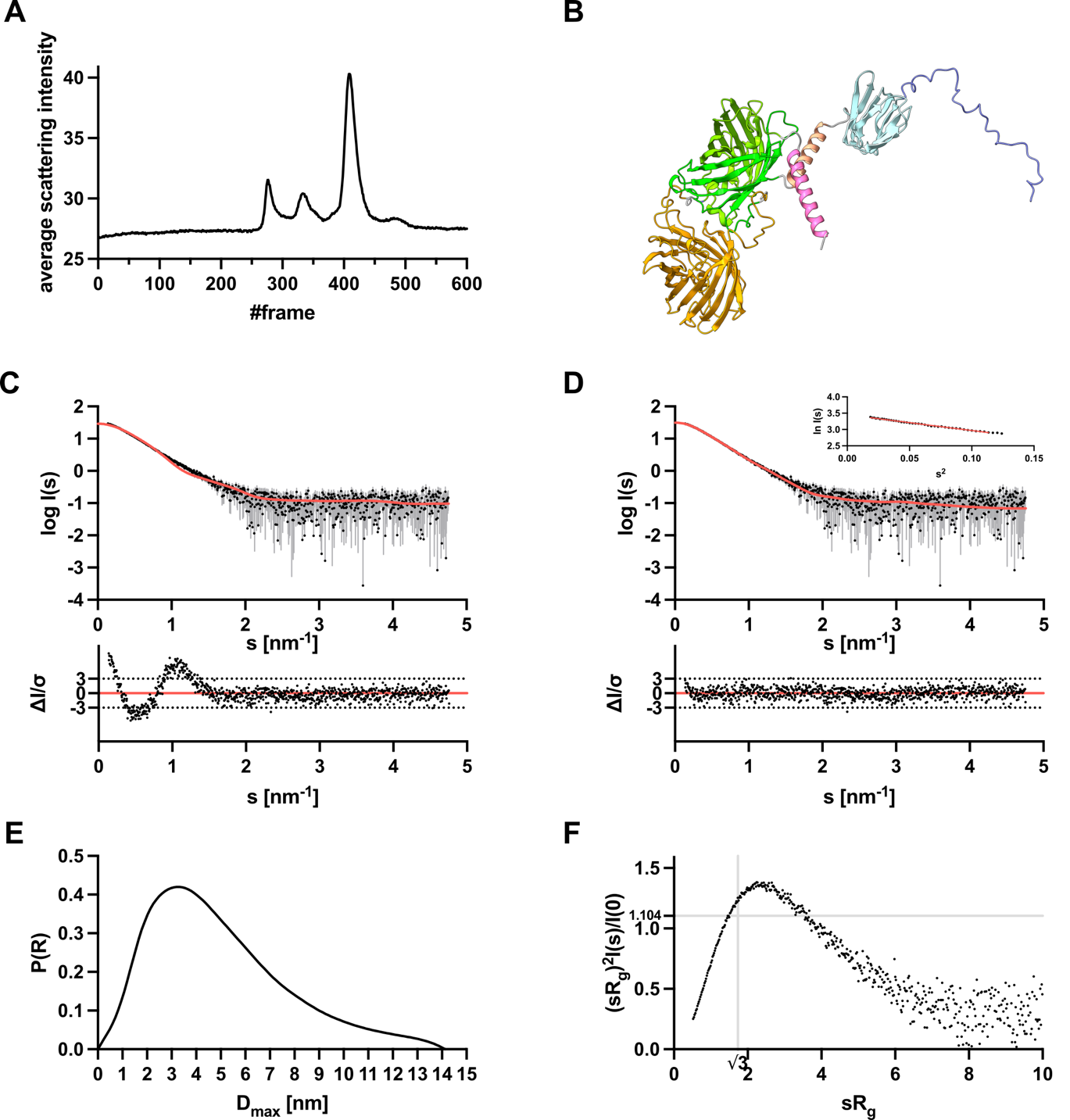

**Figure S4: Small-angle X-ray scattering data from ATPLyzer GO 1.7 m apo. A:** Chromixs SEC SAXS elution profiles. Each frame corresponds to 3 sec. **B**: Final selected RANCH model of ATPLyzer GO 1.7 m. The reporter FP (cpGFP) is shown in green, the reference FP (LSSmOrange) in orange. The FoF1-ATPase ε-subunit (εFoF1) forms the molecular binding element of the biosensor and compose of the N-terminal β-sheet in turquoise, the N-terminal helix in salmon, the C-terminal helix in pink and the N-terminal tail in blue. **C:** Scattering data of ATPLyzer 1.7 m GO apo. Experimental data is shown in black dots, with grey error bars. The theoretical scattering intensity from the initial AlphaFold3 model (*χ* ^2^ value of 5.13), created with CRYSOL, is shown as red line and below is the residual plot of the data. **D:** Experimental data of ATPLyzer 1.7 m GO apo is shown in black dots, with grey error bars. The theoretical scattering intensity from the final selected RANCH model, created with CRYSOL, is shown as red line and below is the residual plot of the data. The Guinier plot is added in the right corner. **E:** *p(r)* function of ATPLyzer 1.7 m GO apo showed an elongated particle. **F:** Dimensionless Kratky plots of ATPLyzer 1.7 m GO apo.

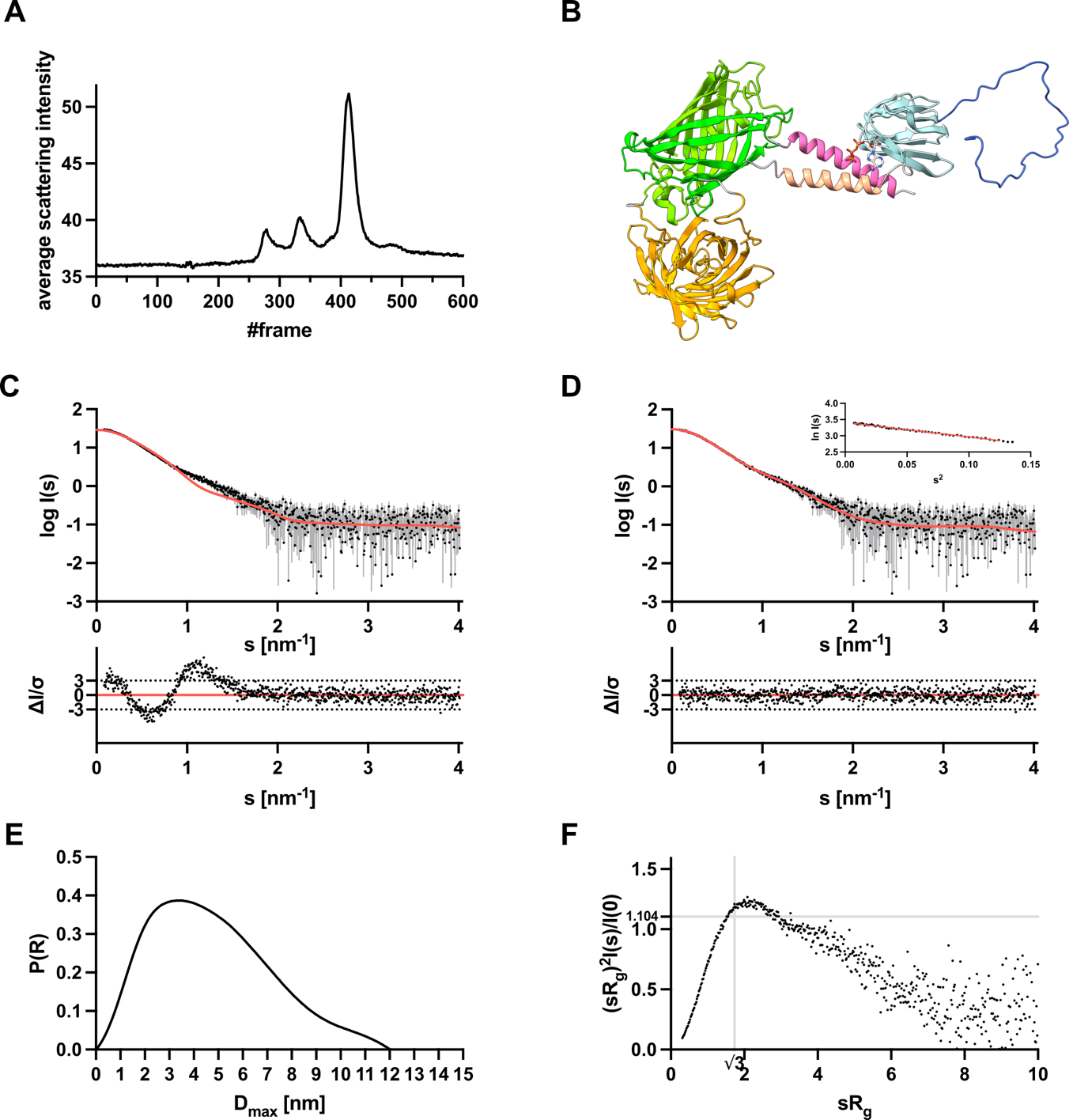

**Figure S5: Small-angle X-ray scattering data from ATPLyzer GO 1.7 m with ATP. A:** Chromixs SEC SAXS elution profiles. Each frame corresponds to 3 sec. **B**: Final selected RANCH model of ATPLyzer GO 1.7 m with ATP docked. The reporter FP (cpGFP) is shown in green, the reference FP (LSSmOrange) in orange. The FoF1-ATPase ε-subunit (εFoF1) forms the molecular binding element of the biosensor and compose of the N-terminal β-sheet in turquoise, the N-terminal helix in salmon, the C-terminal helix in pink and the N-terminal tail in blue. **C:** Scattering data of ATPLyzer 1.7 m GO with ATP. Experimental data is shown in black dots, with grey error bars. The theoretical scattering intensity from the initial AlphaFold3 model (*χ* ^2^ value of 5.35), created with CRYSOL, is shown as red line and below is the residual plot of the data. **D:** Experimental data of ATPLyzer 1.7 m GO with ATP is shown in black dots, with grey error bars. The theoretical scattering intensity from the final selected RANCH model, created with CRYSOL, is shown as red line and below is the residual plot of the data. The Guinier plot is added in the right corner. **E:** *p(r)* function of ATPLyzer 1.7 m GO with ATP showed an elongated particle. **F:** Dimensionless Kratky plots of ATPLyzer 1.7 m GO with ATP.

**Table S 1: Overall SAXS Data**

| **Data collection parameters** | | |
| --- | --- | --- |
| SAXS Device | BM29, ESRF Grenoble ^1^ | |
| Detector | PILATUS 3 X 2M | |
| Detector distance (m) | 2.812 | |
| Beam size | 200 µm x 100 µm | |
| Wavelength (nm) | 0.099 | |
| Sample environment | Quartz glass capillary, 1 mm ø | |
| Absolute scaling method | Comparison with scattering from pure H_2_O | |
| Normalization | To transmitted intensity by beam-stop counter | |
| Scattering intensity scale | Absolute scale, cm^-1^ | |
| *s* range (nm^-1^)^‡^ | 0.025 – 5.5 | |
| **Sample** | **ATPLyzer GO 1.7 m apo** | **ATPLyzer GO 1.7 m with ATP** |
| Organism | *Bacillus spec.* εFoF1 subunit | |
| UniProt ID | P37812, | |
| Mode of measurement | Online SEC-SAXS | |
| Temperature (°C) | 10.0 | |
| SEC-Column | Superdex 200 increase 10/300 | |
| Injection volume (µl) | 100 | |
| Flowrate (ml/min) | 0.6 | |
| Exposure time (# frames) | 3 s (800) | |
| # frames used for averaging | 17 | 10 |
| Protein buffer | 50 mM MOPS pH7.0, 100 mM KCl, 1 mM MgCl_2_ | 50 mM MOPS pH7.0, 100 mM KCl, 1 mM MgCl_2_, 5mM ATP |
| Protein concentration (mg/ml) | 4.9 | 4.9 |
| Substrate | - | 5 mM ATP |
| **Structural parameters** | | |
| *Guinier Analysis (PRIMUS)* | | |
| *I*(0) ± σ (cm^-1^) | 31.56 ± 0.10 | 30.37 ± 0.11 |
| *R*_g_ ± σ (nm) | 3.81 ± 0.02 | 3.67 ± 0.02 |
| *s-range* (nm^-1^) | 0.136 – 0.337 | 0.085 – 0.352 |
| *min < sRg < max limit* | 0.520 – 1.283 | 0.312 – 1.294 |
| Data point range | 1 - 40 | 1 - 53 |
| Linear fit assessment (R^2^) | 0.996 | 0.992 |
| *PDDF/P(r) Analysis (GNOM)* | | |
| *I*(0) ± σ (cm^-1^) | 31.83 ± 0.11 | 30.39 ± 0.10 |
| *R*_g_ ± σ (nm) | 3.99 ± 0.02 | 3.75 ± 0.02 |
| *D*_max_ (nm) | 14.13 | 12.02 |
| Porod volume (nm^3^) | 109.86 | 106.16 |
| *s-range* (nm^-1^) | 0.136 – 4.757 | 0.085 – 4.012 |
| χ2 / CorMap P-value | 1.019 / 0.354 | 1.028 / 0.778 |
| **Molecular mass (kDa)** | | |
| From *I*(0) | n.d. | n.d. |
| From Qp ^2^ | 78.43 | 70.27 |
| From MoW2 ^3^ | 71.13 | 72.22 |
| From Vc ^4^ | 70.81 | 68.85 |
| Bayesian Inference ^5^ | 74.33 | 68.78 |
| From GNNOM ^6^ | 74.00 | 73.10 |
| From sequence | 72.21 (monomer) | |
| **Atomistic modeling** | | |
| CRYSOL (with default parameters, Constant subtraction allowed) | | |
| Structure template | RANCH models (based on AlphaFold3) | |
| *s-*range for fit (nm^-1^) | 0.136 – 4.757 | 0.085 – 4.012 |
| *χ* ^2^, CorMap *P*-value | 1.069 / 0.053 | 0.995 / 0.780 |
| Predicted *R_g_* (nm) | 3.91 | 3.78 |
| **SASBDB accession codes** ^7^ | SASDYQ9 | SASDYR9 |
| **Software** |  | |
| ATSAS Software Version ^8^ | 3.0.5 | |
| Primary data reduction | PRIMUS ^9^ | |
| Data processing | GNOM ^10^ | |
| Model creation | RANCH (part of EOM) ^11, 12^ / AlphaFold3 ^13^ | |
| Structure evaluation | CRYSOL ^14^ | |
| Statistic goodness-of-fit test | χ2, CorMap ^15^ | |
| Model visualization | PyMOL ^16^ | |

‡s = 4πsin(θ)/λ, 2θ – scattering angle, λ – Xray-wavelength, n.d. not determined

**Table S2: List of Primers**

| **Name** | **Sequence (5'-3')** |
| --- | --- |
| Construction of pRSET_B_ 10xHis 1.7 m, 7 µ and n.r variants | |
| Pfw_GFP-C_Ol | GGCATTACACACGGCATG |
| Prev_GFP-N_Ol | CACGCCGGTAAACAGTTC |
| Pfw_QUE_eGFP-C_Ol | CTGTTTACCGGCGTGGTGCCCATCCTGGTC |
| Prev_QUE_eGFP-N_Ol | GCCGTGTGTAATGCCGGCGGCGGTCACGAA |
| Pfw_GFP-C_Ol | GGCATTACACACGGCATG |

**Table S3: Bacterial strains utilized in this study**

| **Strain** | **Relevant characteristics** |
| --- | --- |
| ***Escherichia coli*** |  |
| DH5α^17^ | F^−^ ϕ80*dlac* Δ(*lacZ*)M15 Δ(*lacZYA-argF*) U169 *endA1 recA1 hsdR17* (r_K_^−^ m_K_^+^) *deoR thi-1 phoA supE44* λ^−^ *gyrA96 relA1*; strain used for cloning procedures |
| BL21 (DE3)^18^ | *E. coli* str. B F– *ompT* *gal* *dcm* *lon* *hsdSB* (*rB*–*mB*–) λ (DE3)  [*lacI* *lacUV5*-*T7p07* *ind1* *sam7* *nin5*]) [*malB*+]K-12 (λS); used for biosensor expresion |
| XL1-blue (Agilent Technologies)^19^ | endA1 gyrA96(nal^R^) thi-1 recA1 relA1 lac glnV44 F'[ ::Tn10 proAB^+^ lacI^q^ Δ(lacZ)M15] hsdR17(r_K_^-^ m_K_^+^); used for biosensor basal expresion and in vivo studies |

**Table S4: List of plasmids**

| **Strain** | **Relevant characteristics** |
| --- | --- |
| pRSET-QUE2m^20^,  pRSET-QUE7mu^20^, pRSET-QNA^20^ | Amp, pRSET_B_ derivative for expression of ATPQueen biosensors (cpGFP inserted into the respective insertion sites of ε subunit of *B. subtillis or B. ps3* FoF1 ATPase) with an N-terminal 6x-His-tag, T7-tag and Xpress-tag under the control of T7 promoter, Enterokinase cleavage site |
| pAZ76, pAZ78, pAZ80, pAZ82, pAZ275, pAZ276 | Amp, pRSET_B_ derivative for expression of the matryoshka cassette (cpGFP and nested LSSmOrange LSSmApple inserted into the respective insertion sites of ε subunit of *B. subtillis or B. ps3* FoF1 ATPase) with an N-terminal 6x-His-tag, T7-tag and Xpress-tag under the control of T7 promoter, Enterokinase cleavage site |

**Supplemental Sequences**

>ATPLyzer GA 1.7 m
atgcggggttctcatcatcatcatcatcatggtatggctagcatgactggtggacagcaaatgggtcgggatctgtacgacgatgacgataaggatccgatgaaaactgtgaaagtgaatataacaacccctgatgggccagtctacgacgctgatatcgagatggtgtccgtgcgggccgagagtggtgatctcggcatcctccccggtcacattcccacaaaggccccactgaagatcggagctgtgcggctgaagaaggacggccaaaccgagatggtcgcagtctcaggcggcactgttgaagtgcggcctgaccacgttaccattaatgctcaagccgctgaaacagccgaaggaatcgacaaagagagagcagaagccgcaagacagagggcccaggagcggctgaactctcaaacccgttacaacagccacaacgtctatatcatggccgacaagcagaagaacggcatcaaggtgaacttcaagatccgccacaacatcgaggacggcagcgtgcagctcgccgaccactaccagcagaacacccccatcggcgacggccccgtgctgctgcccgacaaccactacctgagcacccagtccaagctgagcaaagaccccaacgagaagcgcgatcacatggtcctgctggagttcgtgaccgccgccggcattacacacggcatggatgaactgtatggcggcaccgtgagcaagggcgaggagaataacatggccatcatcaaggagttcatgcgcttcaaggtgcacatggagggctccgtgaacggccacgagttcgagatcgagggcgagggcgagggccgcccctacgaggcctttcagaccgctaagctgaaggtgaccaagggtggccccctgcccttcacctgggacatcctgtcccctcagttcatgtacggctccaaggtctacattaagcacccagccgacatccccgactacttcaagctgtccttccccgagggcttcaggtgggagcgcgtgatgatcttcgaggacggcggcattattcacgttaaccaggactcctccctgcaggacggcgtgttcatctacaaggtgaagctgcgcggcaccaacttcccctccgacggccccgttatgcagaagaagaccatgggccttgaggcctgtgaggagcggatgtaccccgaggacggcgccctgaagagcgagtataaggagtggctgaagctgaaggacggcggccactacgccgccgaggtcaagaccacctacaaggccaagaagcccgtgcagctgccaggcgcctacatcgtcgacatcaagttggacatcgtgtcccacaacgaggactacaccatcgtggaacagtacgaacgcgccgagggccgccactccaccggcggcatggacgagctgtacaagggcggcagcgcgagccagggcgaagaactgtttaccggcgtggtgcccatcctggtcgagctggacggcgacgtaaacggccacaagttcagcgtgtccggcgagggcgagggcgatgccacctacggcaagctgaccctgaagttcatctgcaccaccggcaagctgcccgtgccctggcccaccctcgtgaccaccctgacctacggcgtgcagtgcttcagccgctaccccgaccacatgaagcagcacgacttcttcaagtccgccatgcccgaaggctacgtccaggagcgcaccatcttcttcaaggacgacggcaactacaagacccgcgccgaggtgaagttcgagggcgacaccctggtgaaccgcatcgagctgaagggcatcgacttcaaggaggacggcaacatcctggggcacaagctggagtacaacctgggtgacatcgacttcaaacgggccgaactggcgttaaaacgcgccatgaaccgtttgagcgttgcggaaatgaagtga

>ATPLyzer GO 1.7 m
atgcggggttctcatcatcatcatcatcatggtatggctagcatgactggtggacagcaaatgggtcgggatctgtacgacgatgacgataaggatccgatgaaaactgtgaaagtgaatataacaacccctgatgggccagtctacgacgctgatatcgagatggtgtccgtgcgggccgagagtggtgatctcggcatcctccccggtcacattcccacaaaggccccactgaagatcggagctgtgcggctgaagaaggacggccaaaccgagatggtcgcagtctcaggcggcactgttgaagtgcggcctgaccacgttaccattaatgctcaagccgctgaaacagccgaaggaatcgacaaagagagagcagaagccgcaagacagagggcccaggagcggctgaactctcaaacccgttacaacagccacaacgtctatatcatggccgacaagcagaagaacggcatcaaggtgaacttcaagatccgccacaacatcgaggacggcagcgtgcagctcgccgaccactaccagcagaacacccccatcggcgacggccccgtgctgctgcccgacaaccactacctgagcacccagtccaagctgagcaaagaccccaacgagaagcgcgatcacatggtcctgctggagttcgtgaccgccgccggcattacacacggcatggatgaactgtatggcggcaccatggtgagcaagggcgaggagaataacatggccatcatcaaggagttcatgcgcttcaaggtgcgcatggagggctccgtgaacggccacgagttcgagatcgagggcgagggcgagggccgcccctacgagggctttcagaccgttaagctgaaggtgaccaagggtggccccctgcccttcgcctgggacatcttgtcccctcagttcacctacggctccaaggcctacgtgaagcaccccgccgacatccccgactacctcaagctgtccttccccgagggcttcaagtgggagcgcgtgatgaacttcgaggacggcggcgtggtgaccgtgactcaggactcctccctgcaggacggcgagttcatctacaaggtgaagctgcgcggcaccaacttcccctccgacggccccgtaatgcagaagaagaccatgggcatggaggcctcctccgagcggatgtaccccgaggacggcgccctgaagggcgaggacaagctcaggctgaagctgaaggacggcggccactacacctccgaggtcaagaccacctacaaggccaagaagcccgtgcagttgcccggcgcctacatcgtcgacatcaagttggacatcacctcccacaacgaggactacaccatcgtggaacagtacgaacgcgccgagggccgccactccaccggcggcatggacgagctgtacaagggcggcagcgcgagccagggcgaagaactgtttaccggcgtggtgcccatcctggtcgagctggacggcgacgtaaacggccacaagttcagcgtgtccggcgagggcgagggcgatgccacctacggcaagctgaccctgaagttcatctgcaccaccggcaagctgcccgtgccctggcccaccctcgtgaccaccctgacctacggcgtgcagtgcttcagccgctaccccgaccacatgaagcagcacgacttcttcaagtccgccatgcccgaaggctacgtccaggagcgcaccatcttcttcaaggacgacggcaactacaagacccgcgccgaggtgaagttcgagggcgacaccctggtgaaccgcatcgagctgaagggcatcgacttcaaggaggacggcaacatcctggggcacaagctggagtacaacctgggtgacatcgacttcaaacgggccgaactggcgttaaaacgcgccatgaaccgtttgagcgttgcggaaatgaagtga

>ATPLyzer GA 7 µ
atgcggggttctcatcatcatcatcatcatggtatggctagcatgactggtggacagcaaatgggtcgggatctgtacgacgatgacgataaggatccgatgaaaacgatccacgtgagcgtcactactcctgatggcccggtgtacgaagacgatgttgaaatggtcagcgtcaaagcgaaaagcggcgagctcggcattttgccggggcacattccgcttaaggccccgctcgagatcagcgcggcccggctgaaaaaaggcggcaaaacgcaatacattgccgtcagcggcggcaatttggaagtccgcccggacaaagtgacgattaacgctcaagctgctgaacgggcggaggacattgacgtcctccgcgccaaagcggcgaaagagcgggcggagcgccgcctgcaaagccagacccgttacaacagccacaacgtctatatcatggccgacaagcagaagaacggcatcaaggtgaacttcaagatccgccacaacatcgaggacggcagcgtgcagctcgccgaccactaccagcagaacacccccatcggcgacggccccgtgctgctgcccgacaaccactacctgagcacccagtccaagctgagcaaagaccccaacgagaagcgcgatcacatggtcctgctggagttcgtgaccgccgccggcattacacacggcatggatgaactgtatggcggcaccgtgagcaagggcgaggagaataacatggccatcatcaaggagttcatgcgcttcaaggtgcacatggagggctccgtgaacggccacgagttcgagatcgagggcgagggcgagggccgcccctacgaggcctttcagaccgctaagctgaaggtgaccaagggtggccccctgcccttcacctgggacatcctgtcccctcagttcatgtacggctccaaggtctacattaagcacccagccgacatccccgactacttcaagctgtccttccccgagggcttcaggtgggagcgcgtgatgatcttcgaggacggcggcattattcacgttaaccaggactcctccctgcaggacggcgtgttcatctacaaggtgaagctgcgcggcaccaacttcccctccgacggccccgttatgcagaagaagaccatgggccttgaggcctgtgaggagcggatgtaccccgaggacggcgccctgaagagcgagtataaggagtggctgaagctgaaggacggcggccactacgccgccgaggtcaagaccacctacaaggccaagaagcccgtgcagctgccaggcgcctacatcgtcgacatcaagttggacatcgtgtcccacaacgaggactacaccatcgtggaacagtacgaacgcgccgagggccgccactccaccggcggcatggacgagctgtacaagggcggcagcgcgagccagggcgaagaactgtttaccggcgtggtgcccatcctggtcgagctggacggcgacgtaaacggccacaagttcagcgtgtccggcgagggcgagggcgatgccacctacggcaagctgaccctgaagttcatctgcaccaccggcaagctgcccgtgccctggcccaccctcgtgaccaccctgacctacggcgtgcagtgcttcagccgctaccccgaccacatgaagcagcacgacttcttcaagtccgccatgcccgaaggctacgtccaggagcgcaccatcttcttcaaggacgacggcaactacaagacccgcgccgaggtgaagttcgagggcgacaccctggtgaaccgcatcgagctgaagggcatcgacttcaaggaggacggcaacatcctggggcacaagctggagtacaacctgggtgacatcgacttcaaacgggccgaactggcgttaaaacgcgccatgaaccgtttgagcgttgcggaaatgaagtga

>ATPLyzer GO 7 µ
atgcggggttctcatcatcatcatcatcatggtatggctagcatgactggtggacagcaaatgggtcgggatctgtacgacgatgacgataaggatccgatgaaaacgatccacgtgagcgtcactactcctgatggcccggtgtacgaagacgatgttgaaatggtcagcgtcaaagcgaaaagcggcgagctcggcattttgccggggcacattccgcttaaggccccgctcgagatcagcgcggcccggctgaaaaaaggcggcaaaacgcaatacattgccgtcagcggcggcaatttggaagtccgcccggacaaagtgacgattaacgctcaagctgctgaacgggcggaggacattgacgtcctccgcgccaaagcggcgaaagagcgggcggagcgccgcctgcaaagccagacccgttacaacagccacaacgtctatatcatggccgacaagcagaagaacggcatcaaggtgaacttcaagatccgccacaacatcgaggacggcagcgtgcagctcgccgaccactaccagcagaacacccccatcggcgacggccccgtgctgctgcccgacaaccactacctgagcacccagtccaagctgagcaaagaccccaacgagaagcgcgatcacatggtcctgctggagttcgtgaccgccgccggcattacacacggcatggatgaactgtatggcggcaccatggtgagcaagggcgaggagaataacatggccatcatcaaggagttcatgcgcttcaaggtgcgcatggagggctccgtgaacggccacgagttcgagatcgagggcgagggcgagggccgcccctacgagggctttcagaccgttaagctgaaggtgaccaagggtggccccctgcccttcgcctgggacatcttgtcccctcagttcacctacggctccaaggcctacgtgaagcaccccgccgacatccccgactacctcaagctgtccttccccgagggcttcaagtgggagcgcgtgatgaacttcgaggacggcggcgtggtgaccgtgactcaggactcctccctgcaggacggcgagttcatctacaaggtgaagctgcgcggcaccaacttcccctccgacggccccgtaatgcagaagaagaccatgggcatggaggcctcctccgagcggatgtaccccgaggacggcgccctgaagggcgaggacaagctcaggctgaagctgaaggacggcggccactacacctccgaggtcaagaccacctacaaggccaagaagcccgtgcagttgcccggcgcctacatcgtcgacatcaagttggacatcacctcccacaacgaggactacaccatcgtggaacagtacgaacgcgccgagggccgccactccaccggcggcatggacgagctgtacaagggcggcagcgcgagccagggcgaagaactgtttaccggcgtggtgcccatcctggtcgagctggacggcgacgtaaacggccacaagttcagcgtgtccggcgagggcgagggcgatgccacctacggcaagctgaccctgaagttcatctgcaccaccggcaagctgcccgtgccctggcccaccctcgtgaccaccctgacctacggcgtgcagtgcttcagccgctaccccgaccacatgaagcagcacgacttcttcaagtccgccatgcccgaaggctacgtccaggagcgcaccatcttcttcaaggacgacggcaactacaagacccgcgccgaggtgaagttcgagggcgacaccctggtgaaccgcatcgagctgaagggcatcgacttcaaggaggacggcaacatcctggggcacaagctggagtacaacctgggtgacatcgacttcaaacgggccgaactggcgttaaaacgcgccatgaaccgtttgagcgttgcggaaatgaagtga

>ATPLyzer GA n.r.
atgcggggttctcatcatcatcatcatcatggtatggctagcatgactggtggacagcaaatgggtcgggatctgtacgacgatgacgataaggatccgatgaaaactgtgaaagtgaatataacaacccctgatgggccagtctacgacgctgatatcgagatggtgtccgtgcgggccgagagtggtgatctcggcatcctccccggtcacattcccacaaaggccccactgaagatcggagctgtgcggctgaagaaggacggccaaaccgagatggtcgcagtctcaggcggcactgttgaagtgcggcctgaccacgttaccattaatgctcaagccgctgaaacagccgaaggaatcgacaaagagagagcagaagccgcaagacagagggcccaggagcggctgaactctcaaacccgttacaacagccacaacgtctatatcatggccgacaagcagaagaacggcatcaaggtgaacttcaagatccgccacaacatcgaggacggcagcgtgcagctcgccgaccactaccagcagaacacccccatcggcgacggccccgtgctgctgcccgacaaccactacctgagcacccagtccaagctgagcaaagaccccaacgagaagcgcgatcacatggtcctgctggagttcgtgaccgccgccggcattacacacggcatggatgaactgtatggcggcaccgtgagcaagggcgaggagaataacatggccatcatcaaggagttcatgcgcttcaaggtgcacatggagggctccgtgaacggccacgagttcgagatcgagggcgagggcgagggccgcccctacgaggcctttcagaccgctaagctgaaggtgaccaagggtggccccctgcccttcacctgggacatcctgtcccctcagttcatgtacggctccaaggtctacattaagcacccagccgacatccccgactacttcaagctgtccttccccgagggcttcaggtgggagcgcgtgatgatcttcgaggacggcggcattattcacgttaaccaggactcctccctgcaggacggcgtgttcatctacaaggtgaagctgcgcggcaccaacttcccctccgacggccccgttatgcagaagaagaccatgggccttgaggcctgtgaggagcggatgtaccccgaggacggcgccctgaagagcgagtataaggagtggctgaagctgaaggacggcggccactacgccgccgaggtcaagaccacctacaaggccaagaagcccgtgcagctgccaggcgcctacatcgtcgacatcaagttggacatcgtgtcccacaacgaggactacaccatcgtggaacagtacgaacgcgccgagggccgccactccaccggcggcatggacgagctgtacaagggcggcagcgcgagccagggcgaagaactgtttaccggcgtggtgcccatcctggtcgagctggacggcgacgtaaacggccacaagttcagcgtgtccggcgagggcgagggcgatgccacctacggcaagctgaccctgaagttcatctgcaccaccggcaagctgcccgtgccctggcccaccctcgtgaccaccctgacctacggcgtgcagtgcttcagccgctaccccgaccacatgaagcagcacgacttcttcaagtccgccatgcccgaaggctacgtccaggagcgcaccatcttcttcaaggacgacggcaactacaagacccgcgccgaggtgaagttcgagggcgacaccctggtgaaccgcatcgagctgaagggcatcgacttcaaggaggacggcaacatcctggggcacaagctggagtacaacctgggtgacaccgatattcgccgggccgagctggcactgcagaaggccctgaacaagctggacgtggctgggaaggcaaactga

>ATPLyzer GO n.r.
atgcggggttctcatcatcatcatcatcatggtatggctagcatgactggtggacagcaaatgggtcgggatctgtacgacgatgacgataaggatccgatgaaaactgtgaaagtgaatataacaacccctgatgggccagtctacgacgctgatatcgagatggtgtccgtgcgggccgagagtggtgatctcggcatcctccccggtcacattcccacaaaggccccactgaagatcggagctgtgcggctgaagaaggacggccaaaccgagatggtcgcagtctcaggcggcactgttgaagtgcggcctgaccacgttaccattaatgctcaagccgctgaaacagccgaaggaatcgacaaagagagagcagaagccgcaagacagagggcccaggagcggctgaactctcaaacccgttacaacagccacaacgtctatatcatggccgacaagcagaagaacggcatcaaggtgaacttcaagatccgccacaacatcgaggacggcagcgtgcagctcgccgaccactaccagcagaacacccccatcggcgacggccccgtgctgctgcccgacaaccactacctgagcacccagtccaagctgagcaaagaccccaacgagaagcgcgatcacatggtcctgctggagttcgtgaccgccgccggcattacacacggcatggatgaactgtatggcggcaccatggtgagcaagggcgaggagaataacatggccatcatcaaggagttcatgcgcttcaaggtgcgcatggagggctccgtgaacggccacgagttcgagatcgagggcgagggcgagggccgcccctacgagggctttcagaccgttaagctgaaggtgaccaagggtggccccctgcccttcgcctgggacatcttgtcccctcagttcacctacggctccaaggcctacgtgaagcaccccgccgacatccccgactacctcaagctgtccttccccgagggcttcaagtgggagcgcgtgatgaacttcgaggacggcggcgtggtgaccgtgactcaggactcctccctgcaggacggcgagttcatctacaaggtgaagctgcgcggcaccaacttcccctccgacggccccgtaatgcagaagaagaccatgggcatggaggcctcctccgagcggatgtaccccgaggacggcgccctgaagggcgaggacaagctcaggctgaagctgaaggacggcggccactacacctccgaggtcaagaccacctacaaggccaagaagcccgtgcagttgcccggcgcctacatcgtcgacatcaagttggacatcacctcccacaacgaggactacaccatcgtggaacagtacgaacgcgccgagggccgccactccaccggcggcatggacgagctgtacaagggcggcagcgcgagccagggcgaagaactgtttaccggcgtggtgcccatcctggtcgagctggacggcgacgtaaacggccacaagttcagcgtgtccggcgagggcgagggcgatgccacctacggcaagctgaccctgaagttcatctgcaccaccggcaagctgcccgtgccctggcccaccctcgtgaccaccctgacctacggcgtgcagtgcttcagccgctaccccgaccacatgaagcagcacgacttcttcaagtccgccatgcccgaaggctacgtccaggagcgcaccatcttcttcaaggacgacggcaactacaagacccgcgccgaggtgaagttcgagggcgacaccctggtgaaccgcatcgagctgaagggcatcgacttcaaggaggacggcaacatcctggggcacaagctggagtacaacctgggtgacaccgatattcgccgggccgagctggcactgcagaaggccctgaacaagctggacgtggctgggaaggcaaactga

The FASTA sequences are depicted below: ε subunit of FoF1-ATPase (grey), cpGFP (green), LSSmApple (red), LSSmOrange (yellow)

>ATPLyzer GA 1.7 m
MRGSHHHHHHGMASMTGGQQMGRDLYDDDDKDPMKTVKVNITTPDGPVYDADIEMVSVRAESGDLGILPGHIPTKAPLKIGAVRLKKDGQTEMVAVSGGTVEVRPDHVTINAQAAETAEGIDKERAEAARQRAQERLNSQTRYNSHNVYIMADKQKNGIKVNFKIRHNIEDGSVQLADHYQQNTPIGDGPVLLPDNHYLSTQSKLSKDPNEKRDHMVLLEFVTAAGITHGMDELYGGTVSKGEENNMAIIKEFMRFKVHMEGSVNGHEFEIEGEGEGRPYEAFQTAKLKVTKGGPLPFTWDILSPQFMYGSKVYIKHPADIPDYFKLSFPEGFRWERVMIFEDGGIIHVNQDSSLQDGVFIYKVKLRGTNFPSDGPVMQKKTMGLEACEERMYPEDGALKSEYKEWLKLKDGGHYAAEVKTTYKAKKPVQLPGAYIVDIKLDIVSHNEDYTIVEQYERAEGRHSTGGMDELYKGGSASQGEELFTGVVPILVELDGDVNGHKFSVSGEGEGDATYGKLTLKFICTTGKLPVPWPTLVTTLTYGVQCFSRYPDHMKQHDFFKSAMPEGYVQERTIFFKDDGNYKTRAEVKFEGDTLVNRIELKGIDFKEDGNILGHKLEYNLGDIDFKRAELALKRAMNRLSVAEMK*

>ATPLyzer GO 1.7 m
MRGSHHHHHHGMASMTGGQQMGRDLYDDDDKDPMKTVKVNITTPDGPVYDADIEMVSVRAESGDLGILPGHIPTKAPLKIGAVRLKKDGQTEMVAVSGGTVEVRPDHVTINAQAAETAEGIDKERAEAARQRAQERLNSQTRYNSHNVYIMADKQKNGIKVNFKIRHNIEDGSVQLADHYQQNTPIGDGPVLLPDNHYLSTQSKLSKDPNEKRDHMVLLEFVTAAGITHGMDELYGGTMVSKGEENNMAIIKEFMRFKVRMEGSVNGHEFEIEGEGEGRPYEGFQTVKLKVTKGGPLPFAWDILSPQFTYGSKAYVKHPADIPDYLKLSFPEGFKWERVMNFEDGGVVTVTQDSSLQDGEFIYKVKLRGTNFPSDGPVMQKKTMGMEASSERMYPEDGALKGEDKLRLKLKDGGHYTSEVKTTYKAKKPVQLPGAYIVDIKLDITSHNEDYTIVEQYERAEGRHSTGGMDELYKGGSASQGEELFTGVVPILVELDGDVNGHKFSVSGEGEGDATYGKLTLKFICTTGKLPVPWPTLVTTLTYGVQCFSRYPDHMKQHDFFKSAMPEGYVQERTIFFKDDGNYKTRAEVKFEGDTLVNRIELKGIDFKEDGNILGHKLEYNLGDIDFKRAELALKRAMNRLSVAEMK*

>ATPLyzer GA 7 µ
MRGSHHHHHHGMASMTGGQQMGRDLYDDDDKDPMKTIHVSVTTPDGPVYEDDVEMVSVKAKSGELGILPGHIPLKAPLEISAARLKKGGKTQYIAVSGGNLEVRPDKVTINAQAAERAEDIDVLRAKAAKERAERRLQSQTRYNSHNVYIMADKQKNGIKVNFKIRHNIEDGSVQLADHYQQNTPIGDGPVLLPDNHYLSTQSKLSKDPNEKRDHMVLLEFVTAAGITHGMDELYGGTVSKGEENNMAIIKEFMRFKVHMEGSVNGHEFEIEGEGEGRPYEAFQTAKLKVTKGGPLPFTWDILSPQFMYGSKVYIKHPADIPDYFKLSFPEGFRWERVMIFEDGGIIHVNQDSSLQDGVFIYKVKLRGTNFPSDGPVMQKKTMGLEACEERMYPEDGALKSEYKEWLKLKDGGHYAAEVKTTYKAKKPVQLPGAYIVDIKLDIVSHNEDYTIVEQYERAEGRHSTGGMDELYKGGSASQGEELFTGVVPILVELDGDVNGHKFSVSGEGEGDATYGKLTLKFICTTGKLPVPWPTLVTTLTYGVQCFSRYPDHMKQHDFFKSAMPEGYVQERTIFFKDDGNYKTRAEVKFEGDTLVNRIELKGIDFKEDGNILGHKLEYNLGDIDFKRAELALKRAMNRLSVAEMK*

>ATPLyzer GO 7 µ
MRGSHHHHHHGMASMTGGQQMGRDLYDDDDKDPMKTIHVSVTTPDGPVYEDDVEMVSVKAKSGELGILPGHIPLKAPLEISAARLKKGGKTQYIAVSGGNLEVRPDKVTINAQAAERAEDIDVLRAKAAKERAERRLQSQTRYNSHNVYIMADKQKNGIKVNFKIRHNIEDGSVQLADHYQQNTPIGDGPVLLPDNHYLSTQSKLSKDPNEKRDHMVLLEFVTAAGITHGMDELYGGTMVSKGEENNMAIIKEFMRFKVRMEGSVNGHEFEIEGEGEGRPYEGFQTVKLKVTKGGPLPFAWDILSPQFTYGSKAYVKHPADIPDYLKLSFPEGFKWERVMNFEDGGVVTVTQDSSLQDGEFIYKVKLRGTNFPSDGPVMQKKTMGMEASSERMYPEDGALKGEDKLRLKLKDGGHYTSEVKTTYKAKKPVQLPGAYIVDIKLDITSHNEDYTIVEQYERAEGRHSTGGMDELYKGGSASQGEELFTGVVPILVELDGDVNGHKFSVSGEGEGDATYGKLTLKFICTTGKLPVPWPTLVTTLTYGVQCFSRYPDHMKQHDFFKSAMPEGYVQERTIFFKDDGNYKTRAEVKFEGDTLVNRIELKGIDFKEDGNILGHKLEYNLGDIDFKRAELALKRAMNRLSVAEMK*

>ATPLyzer GA n.r.
MRGSHHHHHHGMASMTGGQQMGRDLYDDDDKDPMKTVKVNITTPDGPVYDADIEMVSVRAESGDLGILPGHIPTKAPLKIGAVRLKKDGQTEMVAVSGGTVEVRPDHVTINAQAAETAEGIDKERAEAARQRAQERLNSQTRYNSHNVYIMADKQKNGIKVNFKIRHNIEDGSVQLADHYQQNTPIGDGPVLLPDNHYLSTQSKLSKDPNEKRDHMVLLEFVTAAGITHGMDELYGGTVSKGEENNMAIIKEFMRFKVHMEGSVNGHEFEIEGEGEGRPYEAFQTAKLKVTKGGPLPFTWDILSPQFMYGSKVYIKHPADIPDYFKLSFPEGFRWERVMIFEDGGIIHVNQDSSLQDGVFIYKVKLRGTNFPSDGPVMQKKTMGLEACEERMYPEDGALKSEYKEWLKLKDGGHYAAEVKTTYKAKKPVQLPGAYIVDIKLDIVSHNEDYTIVEQYERAEGRHSTGGMDELYKGGSASQGEELFTGVVPILVELDGDVNGHKFSVSGEGEGDATYGKLTLKFICTTGKLPVPWPTLVTTLTYGVQCFSRYPDHMKQHDFFKSAMPEGYVQERTIFFKDDGNYKTRAEVKFEGDTLVNRIELKGIDFKEDGNILGHKLEYNLGDTDIRRAELALQKALNKLDVAGKAN*

>ATPLyzer GO n.r.
MRGSHHHHHHGMASMTGGQQMGRDLYDDDDKDPMKTVKVNITTPDGPVYDADIEMVSVRAESGDLGILPGHIPTKAPLKIGAVRLKKDGQTEMVAVSGGTVEVRPDHVTINAQAAETAEGIDKERAEAARQRAQERLNSQTRYNSHNVYIMADKQKNGIKVNFKIRHNIEDGSVQLADHYQQNTPIGDGPVLLPDNHYLSTQSKLSKDPNEKRDHMVLLEFVTAAGITHGMDELYGGTMVSKGEENNMAIIKEFMRFKVRMEGSVNGHEFEIEGEGEGRPYEGFQTVKLKVTKGGPLPFAWDILSPQFTYGSKAYVKHPADIPDYLKLSFPEGFKWERVMNFEDGGVVTVTQDSSLQDGEFIYKVKLRGTNFPSDGPVMQKKTMGMEASSERMYPEDGALKGEDKLRLKLKDGGHYTSEVKTTYKAKKPVQLPGAYIVDIKLDITSHNEDYTIVEQYERAEGRHSTGGMDELYKGGSASQGEELFTGVVPILVELDGDVNGHKFSVSGEGEGDATYGKLTLKFICTTGKLPVPWPTLVTTLTYGVQCFSRYPDHMKQHDFFKSAMPEGYVQERTIFFKDDGNYKTRAEVKFEGDTLVNRIELKGIDFKEDGNILGHKLEYNLGDTDIRRAELALQKALNKLDVAGKAN*
